# CSNAP, the smallest CSN subunit, modulates proteostasis through cullin-RING ubiquitin ligases

**DOI:** 10.1101/433532

**Authors:** Maria G. Füzesi-Levi, Radoslav Ivanov Enchev, Gili Ben-Nissan, Yishai Levin, Meital Kupervaser, Gilgi Friedlander, Tomer Meir Salame, Reinat Nevo, Matthias Peter, Michal Sharon

**Author notes:** Corresponding author: Michal Sharon.

## Abstract

The cullin-RING ubiquitin E3 ligase (CRL) family consists of ~250 complexes that catalyze ubiquitylation of proteins to achieve cellular regulation. All CRLs are inhibited by the COP9 signalosome complex (CSN) through both enzymatic (deneddylation) and non-enzymatic (steric) mechanisms. The relative contribution of these two mechanisms is unclear. Here, we decouple the mechanisms using CSNAP, the recently discovered ninth subunit of the CSN. We find that CSNAP reduces the affinity of CSN toward CRL complexes. Removing CSNAP does not affect deneddylation, but leads to global effects on the CRL, causing altered reproductive capacity, suppressed DNA damage response, decreased viability, and delayed cell cycle progression. Thus, although CSNAP is only 2% of the CSN mass, it plays a critical role in the steric regulation of CRLs by the CSN.

## Introduction

Protein degradation is one of the essential mechanisms that enables reshaping of the proteome landscape in response to various stimuli (Hershko, Ciechanover et al., 2000). The specificity of this process is largely mediated by E3 ligases that ubiquitinate target proteins (Deshaies & Joazeiro, 2009, Enchev, Schulman et al., 2015). One of the largest E3 ubiquitin ligase families, responsible for ubiquitination of 20% of the proteins degraded by the 26S proteasome, is comprised of cullin-RING ligases (CRLs) (Soucy, Smith et al., 2009). This family encompasses ~ 250 distinct complexes that are built in a modular fashion around a central cullin scaffold, which is associated with a specific substrate receptor, adaptor protein, and a RING protein that recruits the E2 enzyme (reviewed in (Deshaies & Joazeiro, 2009, Skaar, Pagan et al., 2013)). Seven different cullins have been identified in humans, each interacting with a dedicated set of receptors, forming CRL complexes that target a single or a small group of substrate proteins. At any given time, various CRLs are active, and their dynamic assembly and disassembly enables cellular adaptation in response to regulatory inputs.

In spite of the great diversity of CRLs in terms of composition and substrate specificity, all complexes are regulated by the COP9 signalosome complex (CSN) (Deshaies & Joazeiro, 2009). The CSN regulates CRLs by means of two independent mechanisms, catalytic and non-catalytic. The first involves enzymatic deconjugation of the ubiquitin-like protein Nedd8 from the cullin subunit (deneddylation) (Cope, Suh et al., 2002). The latter is mediated through physical binding to CRLs, sterically precluding interactions with E2 enzymes and ubiquitination of substrates (Emberley, Mosadeghi et al., 2012, Enchev, Scott et al., 2012, Fischer, Scrima et al., 2011). By inhibiting CRL activity, both mechanisms control the gateway to the exchange cycle that remodels CRL composition (Liu, Reitsma et al., 2018, Mosadeghi, Reichermeier et al., 2016, Reitsma, Liu et al., 2017).

The CSN is a highly conserved complex that exists in all eukaryotes (Wei & Deng, 2003, Wei, Serino et al., 2008). Three types of subunits constitute this complex: two MPN subunits (for Mpr1p and Pad1p N terminal) CSN5 and CSN6 (Glickman, Rubin et al., 1998), six PCI subunits (for proteasome, COP9, and initiation factor 3); CSN1-CSN4, CSN7, and CSN8 (Hofmann & Bucher, 1998); and an additional small, non-PCI or MPN subunit that we recently discovered and termed CSNAP, for CSN acidic protein (Rozen, Fuzesi-Levi et al., 2015). The CSNAP protein consists of only 57 amino acids (molecular weight: 6.2 kDa) that link together the two distinct structural elements of the CSN by mutually binding the MPN subunits CSN5 and CSN6, and the PCI subunit CSN3 (Rozen et al., 2015). Given the small size of CSNAP, a natural question that arises is whether it is actually crucial for CSN function and, if so, what is its functional role? In this study, we address these questions by combining biochemical and cell biology approaches, together with mass spectrometry analysis.

Using the above approaches, we discovered that manipulating CSNAP enables us to uncouple the steric and catalytic activities of the CSN complex. Although it is only 2% of the CSN mass, we find that removing CSNAP has a global effect on the cell cycle, cell viability and DNA damage response. This effect is due to a reduction in the K_d_ of CSN-CRL binding, leaving deneddylation activity unchanged. These findings provides a role for CSNAP, and points to the affinity of CSN-CRL interactions as a critical component for proteostasis.

## Results

### CSNAP alters the strength of CSN-CRL interaction

To investigate the impact of CSNAP on both the enzymatic and steric activities of CSN, we initially examined the complex’s deneddylation activity, using HAP-1 cell lines lacking CSNAP (ΔCSNAP cells) (Rozen et al., 2015). Comparison of the deneddylated/neddylated ratio between WT and ΔCSNAP cells showed that in the absence of CSNAP there are only minor changes of less than 15% of the cullin’s deneddylated fraction (Fig 1A). This result is in accordance with our previous finding, showing that WT and ΔCSNAP cells exhibit a similar rate of deneddylation (Rozen et al., 2015) and with a study that compared the rate of deneddylation of endogenous CSN prepared from HEK293 cells, with that of recombinant CSN^ΔCSNAP^ expressed in insect cells (Emberley et al., 2012, Enchev et al., 2012). Thus, it can be concluded that CSNAP does not significantly affect the catalytic capacity of the CSN complex.

**Figure 1.**
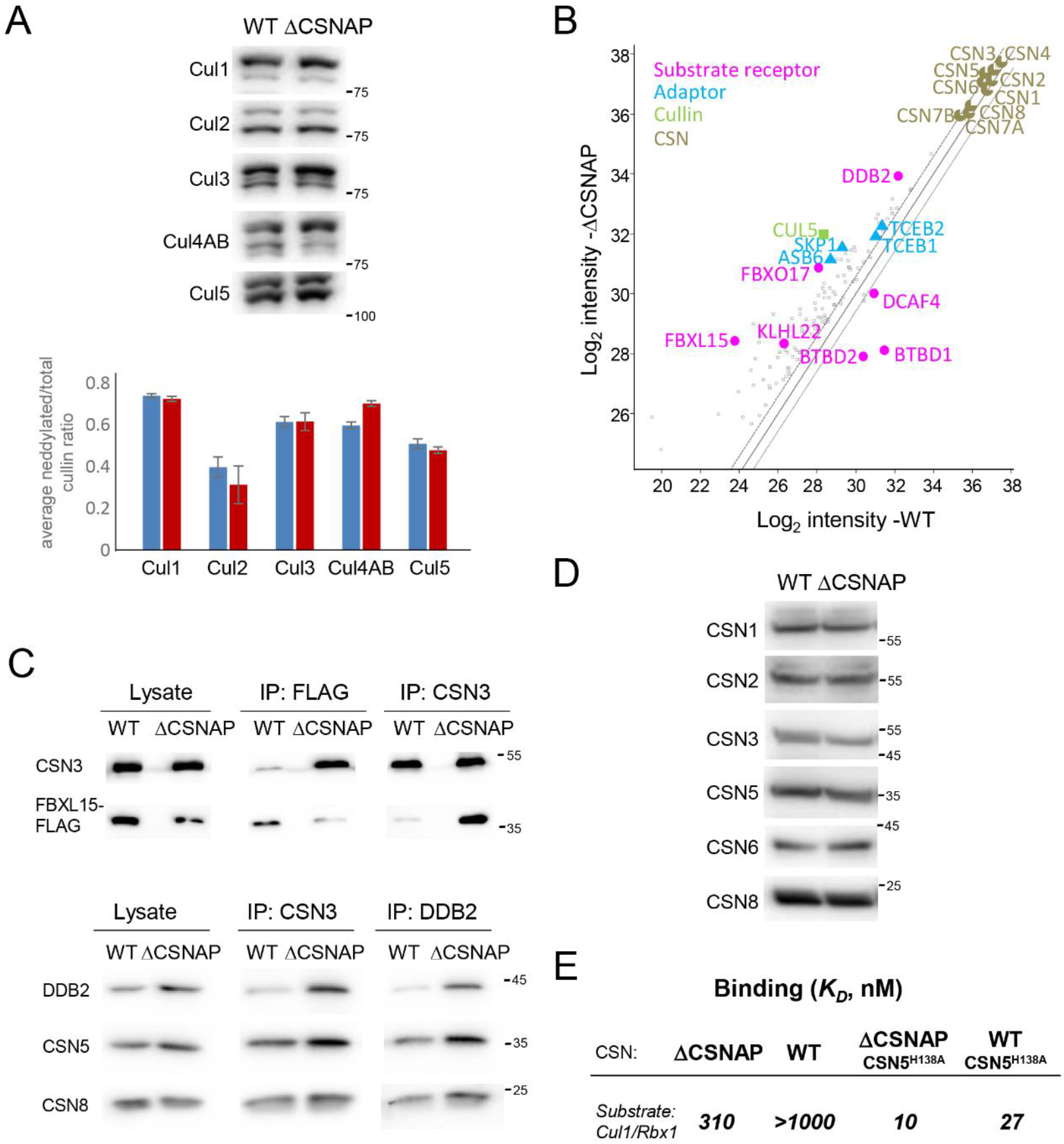
CSNAP modifies the strength of CSN-CRL interactions. (A) CSN catalytic activity is not significantly affected by the absence of CSNAP. A representative western blot of WT and ΔCSNAP cell extracts visualized using antibodies against various cullins (top) and a plot demonstrating the average deneddylated fraction. The graph represents the averages of three independent experiments, with standard errors. (B) CSN and its interacting proteins were pulled down using an antibody against CSN3 from WT and ΔCSNAP cells. Immunoprecipitated proteins were then analyzed by label-free proteomics approach using three biological replicates. Scatter plot comparing log_2_ intensities of proteins in ΔCSNAP and WT samples show that a number of CRL proteins were found to be over- or underrepresented in the pulldown of the CSN and CSN^ΔCSNAP^ complexes. In contrast, the ratio of average intensities for CSN subunits did not exceed the fold change of ΔCSNAP/WT > 1.5, which was considered to be the cut-off for fold change. (C) Validation of the proteomics data for FBXL15/CSN and DDB2/CSN interactions. Reciprocal immunoprecipitation shows a tighter CSN3 interaction with FBXL15 and DDB2, in the absence of CSNAP. (D) The levels of CSN subunits are comparable in WT and ΔCSNAP cells; thus, the differences in the amount of the pulled-down proteins are likely due to different interaction affinities. Representative blot out of three repeats. (E) Determination of the dissociation constant (Kd) for the CSN and CSN^ΔCSNAP^ complexes, and dansyl-labeled Cul1-N8/Rbx1. The absence of CSNAP causes tighter binding to cullin1/Rbx1.

To examine whether the steric activity of CSN^ΔCSNAP^ is affected by the absence of CSNAP, we applied label-free quantification of protein intensities from pull-down assays, coupled with mass spectrometry (MS) analysis of WT and ΔCSNAP cells. We reasoned that if CSNAP impacts the CSN-CRL interaction, differences in the array of protein binding partners will be revealed. Our results indicated that multiple CRL components are significantly enriched in ΔCSNAP pulldowns, in comparison to immunoprecipitation of WT cells (Fig. 1B and Table S1). These mainly include substrate receptors (DDB2, FBOX17, FBXL15 and KLH22) and adapter proteins (TCEB2, TCEB1, SKP1 and ASB6). In the WT cells, only three proteins, DCAF4, BTBD2 and BTBD1, were enriched; all are CRL substrate receptor proteins. To validate these results, we carried out reciprocal co-immunoprecipitation experiments. The results obtained for the ΔCSNAP and WT cell lines confirmed that FBXL15 and DDB2 are enriched in the CSN^ΔCSNAP^ pulldown experiment, in comparison to the WT complex (Fig. 1C). Notably, these results did not arise from changes in the expression levels of CSN subunits, as all CSN subunits (except for CSNAP, depleted from the cells) displayed insignificant differences when the two cell lines were compared (Fig. 1B), a finding that was further validated by Western blot analysis (Fig. 1D, Fig. S1). Taken together, the data suggest that CSNAP plays a role in tuning CSN-CRL interactions in cells.

To further assess the contribution of CSNAP to the CSN/CRL interaction, we utilized a quantitative *in vitro* binding assay to determine the affinity between Cul1-Rbx1, and recombinant CSN^ΔCSNAP^ or CSN complexes (Mosadeghi et al., 2016) (Fig. S2, S3). In this assay, the environmentally-sensitive dye dansyl was conjugated to the C-terminus of Cul1, and an increase in fluorescence upon CSN binding was detected (Mosadeghi et al., 2016). Both WT CSN5 and the well characterized CSN5-H138A mutant (Emberley et al., 2012, Enchev et al., 2012, Mosadeghi et al., 2016) (CSN^5H138A^) were used, as the latter binds Cul1-Rbx1 ~30-fold more tightly, enabling us to reach saturation. The results indicate that CSN complexes display decreased affinity to Cul1-Rbx1, in comparison to CSN^ΔCSNAP^ (Fig. 1E, Fig S4). Taken together, the results imply that CSNAP reduces the affinity of the CSN towards CRL complexes.

### CSNAP is required for proper cell cycle progression and viability

The apparent difference in *K_d_* for CSN^ΔCSNAP^ binding to SCF, compared to CSN, led us to question whether such a change in affinity can influence the repertoire of active CRLs and, as a consequence, the array of ubiquitinated proteins. We therefore performed a large-scale analysis of protein ubiquitination, relying on the enrichment of ubiquitinated tryptic peptides (Udeshi, Mertins et al., 2013). The relative differences in ubiquitination of WT and ΔCSNAP cells was quantified, using the SILAC (stable isotope labeling by amino acids in cell culture) approach (Ong, Blagoev et al., 2002). The results indicated that differences exist in the extent of ubiquitination in ΔCSNAP and WT cell lines (Fig. 2A and Table S2A and B). In all, 159 ubiquitinated proteins, were found to be enriched only in WT and not in ΔCSNAP cells, while 79 ubiquitinated proteins, were abundant in cells lacking CSNAP. Of the 238 proteins whose ubiquitination levels differed between the two type of cells, 64 are known substrates of the different CRL complexes (Table S3) (Emanuele, Elia et al., 2011, Koren, Timms et al., 2018, Yen & Elledge, 2008, Zheng, Zhou et al., 2016). These results suggest that reducing the affinity between CSN and CRL modifies CRL assembly and, consequently, the repertoire of ubiquitinated proteins.

**Figure 2.**
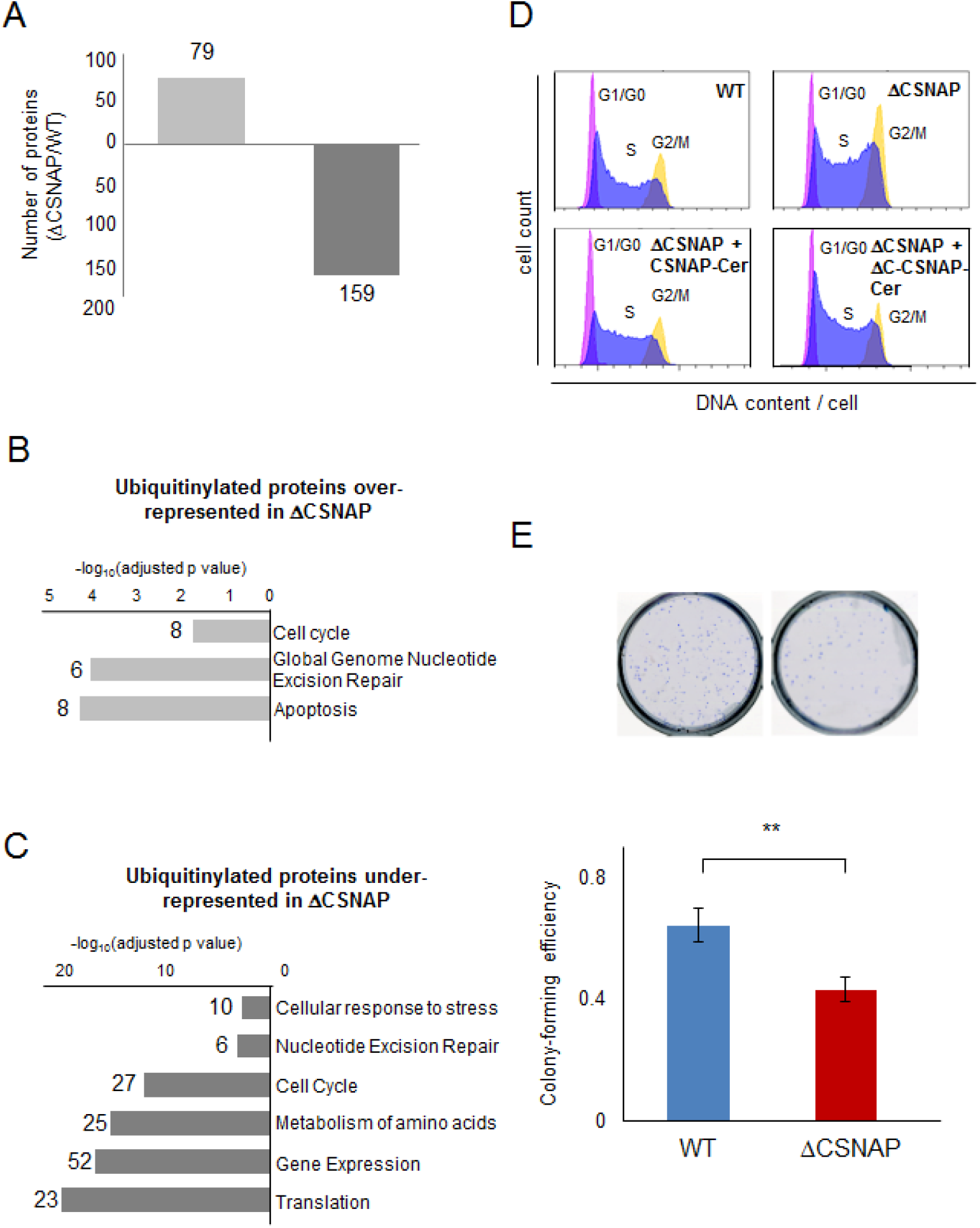
The absence of CSNAP impairs cell cycle progression and viability. (A) We monitored protein ubiquitinylation in stable heavy and light isotope labelled ΔCSNAP and WT cells. The bar plot shows the number of differentially ubiquitinated (up or down) proteins that appeared in at least two out of four independent experiments. (B-C) Pathway analysis of the proteins that are differentially ubiqiutinylated in ΔCSNAP and WT cells. (D) Validation of the influence of the absence of CSNAP on cell cycle. Histograms of BrdU and propidium iodide stained asynchronous cells show that the lack of CSNAP results in a S-G2 shifted phenotype, that can be rescued by the expression of CSNAP-Cerulean, but not when its C-terminal CSN interacting domain is absent. The figure shows a representative experiment out of three. (E) Cells lacking CSNAP have lower colony forming potential than WT cells. The graph represents the results from 14 biological replicates, significance was calculated using Student’s t-test (** p<0.001). Images above show a representative experiment.

Functional annotations revealed that among the ubiquitinated proteins identified as being enriched in WT, or ΔCSNAP cells, 16% are clustered in the cell cycle pathway (Fig. 2B-C, Tables S4, S5A and S5B). We confirmed this data by assessing the cell cycle distribution of both ΔCSNAP and WT cells, using flow cytometry analysis. The results indicated that compared to WT cells, ΔCSNAP cells display larger S and G2 phase populations (Fig. 2D). This phenotype can be prevented by exogenous expression of CSNAP-Cerulean, but not by the truncated form of the protein (ΔC-CSNAP-Cerulean), which lacks the C-terminal region that is crucial for the protein’s integration into the CSN (Rozen et al., 2015). In addition, colony formation assays (Franken, Rodermond et al., 2006) showed that the viability of cells lacking CSNAP is significantly reduced, in comparison to that of WT cells (Fig. 2E). Taken together, our results suggest that the absence of CSNAP influences CSN-CRL interactions in a manner that affects, protein ubiquitination and, therefore likely, cell cycle coordination.

### Cellular protein levels are influenced by CSNAP

Considering the dependence of the ubiquitinated proteome on the presence of CSNAP, we wished to examine whether the impact of this subunit would also be detected in a global proteome analysis. To this end, we performed label-free quantification (Shalit, Elinger et al., 2015) of the proteomes of WT and ΔCSNAP cells. Given that the CSN complex and protein ubiquitination are vital to the DNA damage response (Dubois, Gerber et al., 2016, Fuzesi-Levi, Ben-Nissan et al., 2014, Hannss & Dubiel, 2011, Meir, Galanty et al., 2015), we performed the analysis both prior to and following exposure of the cells to UV irradiation. Data were analyzed by two-way ANOVA, taking into consideration both the UV treatment, and the type of cell being treated (WT or ΔCSNAP). Proteins that were considered significantly differentially expressed were clustered into five groups according to their cellular functions (Fig. 3A and Tables S6A-D and S7). Remarkably, we noticed that cellular pathways that were enriched in this experiment are in accordance with those identified in the SILAC-based ubiquitinylation analysis (Fig. 2B-C). Among these proteins we could identify known substrates of various CRL complexes (Table S8), which are known to be involved in ubiquitination, apoptosis, cell cycle regulation and DNA damage response. This observation may explain the detected phenotypic effects.

**Figure 3.**
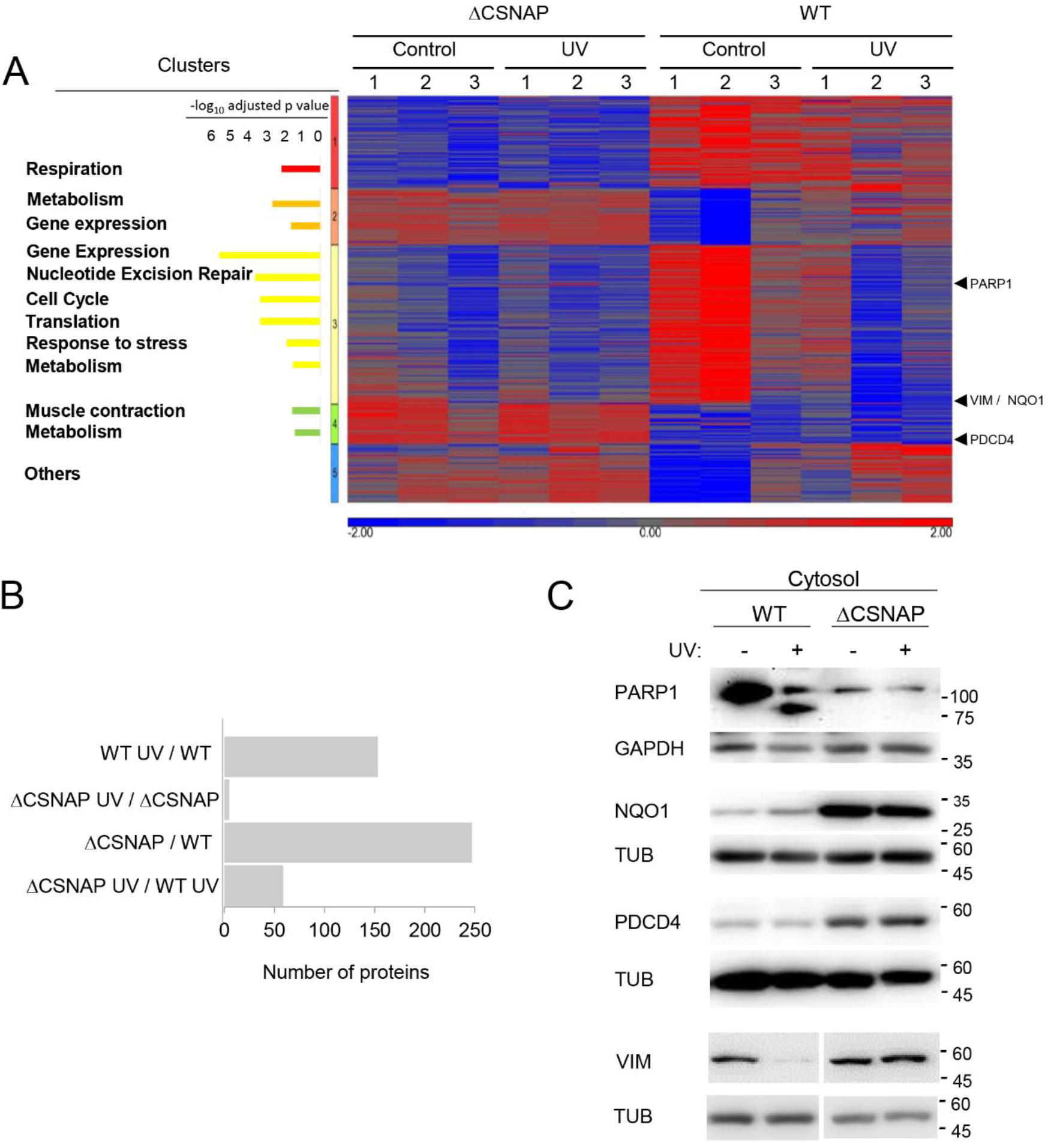
The absence of CSNAP affects proteome remodeling following UV damage. (A) Proteomes of untreated or UV exposed WT and ΔCSNAP cells four hours post damage were analyzed using label free proteomics approach. Proteomics data of three biological replicates, after logarithmic transformation and flooring, were analyzed by two-way ANOVA using the two factors: strain and UV treatment, as well as their interaction. Proteins with p-value below 0.05 and an absolute fold change above 1.5 were considered as being differentially expressed. Heatmap of differentially expressed proteins grouped to five clusters. Pathway analysis of the clusters indicate up- and downregulation of several cellular functions in ΔCSNAP cells. The bar chart on the left expresses the significance levels of the enrichment analysis of the proteins using the protein coding part of the human genome. (B) Comparison of the differentially expressed proteins in the proteome in untreated and UV-exposed WT and ΔCSNAP cells. The bar plot shows each of the four pair-wise comparisons, highlighting that in ΔCSNAP cells the DNA damage response is compromised. (C) Expression levels of four representative proteins: PARP1, NQO1, PDCD4, and vimentin, analyzed by Western blots of WT and ΔCSNAP cell lysates.

Examination of the five clusters indicated that even under normal conditions, there are clear differences in protein expression levels among WT and ΔCSNAP cells. A particularly striking observation was that the cellular response to UV irradiation was nearly abolished in ΔCSNAP cells (Fig. 3B). We validated the proteomics results by Western blot analysis to monitor the expression levels of four proteins that displayed differential expression levels between WT and ΔCSNAP cells: the quinone reductase enzyme, NQO1, the tumor suppressor PDCD4, and the filament protein vimentin, which appear in Cluster 4, and PARP1, a member of the PARP family that appears in Cluster 3 (Fig. 3A; see arrows on the right). The results confirmed that unlike WT cells, NQO1, PDCD4, and vimentin are expressed at high levels in ΔCSNAP cells (Fig. 3C and S5). Likewise, the blots validated that the expression of PARP1 in WT cells is high under normal conditions, and is decreased following UV-induced DNA damage; while in ΔCSNAP cells, regardless of UV irradiation, low levels of PARP1 expression are maintained (Fig. 3C and S5). We also noticed that in WT cells, a cleavage product of PARP1 was detected, a phenomenon that was not observed in ΔCSNAP cells; this result will be further discussed in the next section. In summary, our findings suggest that the lack of CSNAP following UV-treatment elicited a strong and specific influence on downstream effectors of the DNA damage response.

### CSNAP is required for DNA repair

Building on our results reflecting the compromised protein remodeling capability following DNA damage in ΔCSNAP cells, we wished to explore the DNA repair response in these cells. Initially, we measured the DNA repair capacity following UV irradiation using the comet assay (Olive & Banath, 2006). The results indicated that ΔCSNAP cells display a longer tail moment, which is associated with the accumulation of both single- and double-strand DNA breaks (Fig. 4A). Following DNA damage, cells would reduce their rates of proliferation, in order to enable DNA damage repair (Gentile, Latonen et al., 2003). We therefore measured cell proliferation before and after treatment with UV irradiation, and found that, as expected, proliferation arrest was detected in WT cells, however, not in cells lacking CSNAP, nor in ΔCSNAP cells exogenously expressing AC-CSNAP-Cerulean, which is not incorporated into the CSN complex (Fig. 4B). Nevertheless, cell rescue was achieved in ΔCSNAP cells by overexpressing the full-length CSNAP protein.

**Figure 4.**
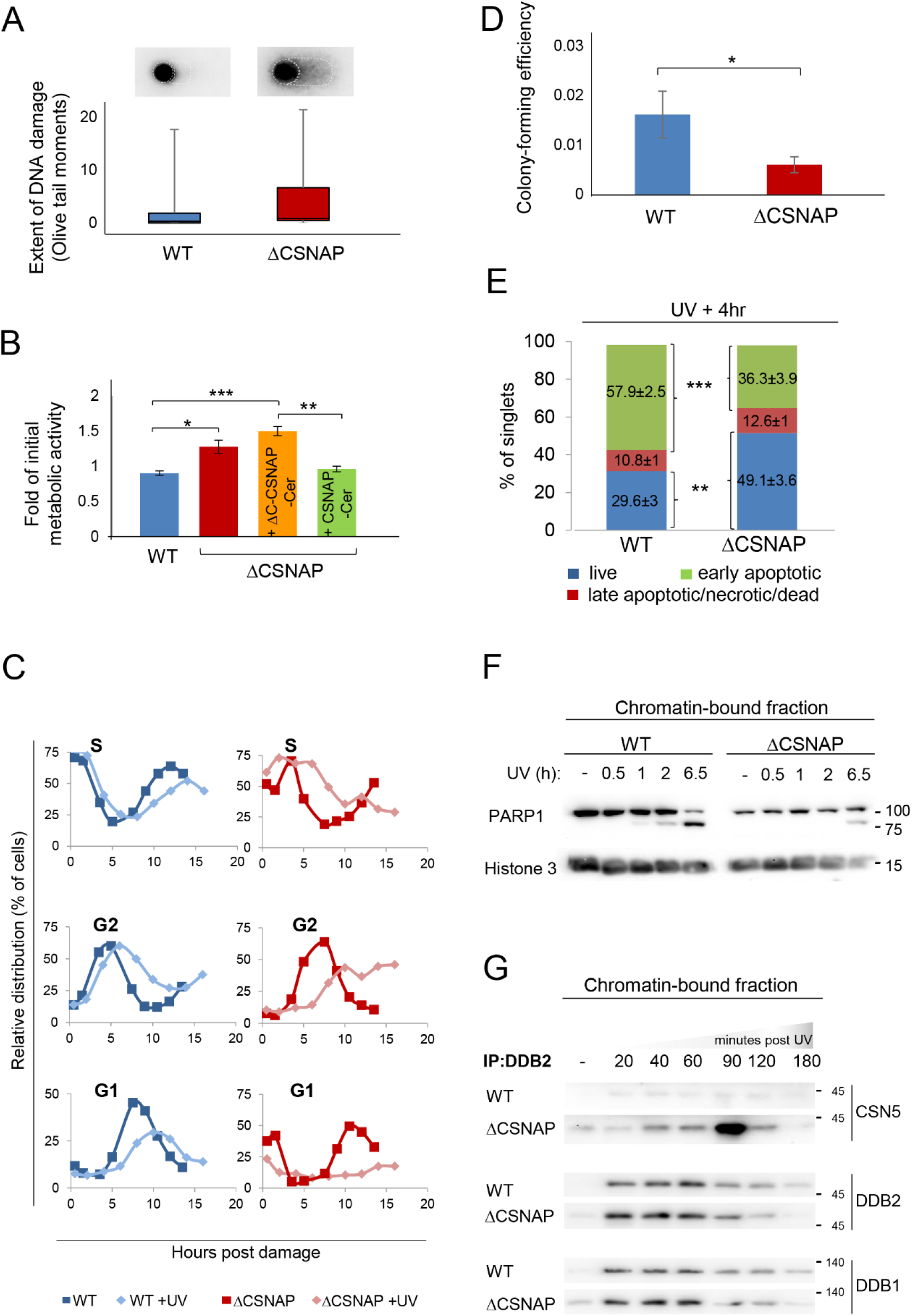
The absence of CSNAP compromises the DNA damage response. (A) Unlike WT cells, damaged DNA accumulates in ΔCSNAP cells following UV exposure. The genotoxic effect of UV was measured using an alkaline comet assay. DNA damage, expressed as Olive tail moments were calculated and presented as a box-whisker plot. Images above the plot display a representative comet shape for each cell type. (B) ΔCSNAP cells fail to down-regulate their metabolic activity after UV-induced DNA damage. The plot shows metabolic activities of WT and ΔCSNAP cells two hours after exposure to UV irradiation, calculated as a fold of initial activity for each cell line. UV-induced DNA damage caused a significant reduction of metabolic activity in WT cells (blue), while cells deficient in CSNAP (red) failed to down-regulate their metabolic activity to that extent. This UV-response phenotype is rescued by exogenous expression of the full-length CSNAP protein (green), but not when its C-terminal CSN-interaction domain was missing (orange). The graph represents the averages of three independent experiments, with standard errors. Significance was calculated using a 2-way ANOVA test, followed by a Tukey Post-Hoc Test (p<0.005). (*p<0.05; **p<0.01; ***p<0.005). (C) UV-exposed ΔCSNAP cells stay longer in S and G2 phases. Comparison of the relative distribution of cell populations in different phases of the cell cycle, as calculated from flow cytometry histograms of double thymidine-synchronized cells, with or without exposure to UV irradiation. UV-caused DNA damage elongates the S and G2 phases; this effect is significantly more pronounced in cells lacking CSNAP. (D) Cells lacking CSNAP exhibit a compromised recovery after exposure to high-dose UV. WT and ΔCSNAP cells were exposed to UV irradiation, prior to incubation in culturing conditions for 8 days. Colonies were stained and counted. ΔCSNAP cells exhibit ~2.7-fold less colony-forming potential following UV damage, in comparison to WT cells. The graph represents average results from 7 biological replicates with standard errors. Significance was calculated using a Student’s t-test (p<0.05). (E) The early apoptotic response is delayed in ΔCSNAP cells, following UV damage. The bar charts represents the percentage of live, early and late apoptotic cells detected by flow cytometry of 6 independent experiments. A significant difference is seen in the percentage of live and early apoptotic cell populations between WT and ΔCSNAP cells, after exposure to UV (***p<0.0005 and **p<0.001, respectively). (F) PARP1 cleavage is delayed in cells lacking CSNAP. Chromatin-bound fractions were monitored by Western blot for caspase-mediated PARP1 cleavage, a marker for commitment to apoptosis. (G) CSN^ΔCSNAP^ exhibits increased affinity towards DDB2, in comparison to the CSN complex. WT and ΔCSNAP cells were exposed to UV irradiation, and DDB2 was immunoprecipitated from the chromatin-bound fraction at different time points post-UV damage. Western blot analyses show tighter CSN-CRL binding when CSNAP is absent. Representative blot out of four repeats.

Considering that widespread DNA damage induces cell cycle arrest (Gentile et al., 2003), we evaluated the cell cycle distribution of WT and ΔCSNAP cells exposed to UV irradiation following a double thymidine block, which induces a G1/S-phase arrest. After their release from cell cycle synchronization, untreated ΔCSNAP cells proceeded to the S phase significantly more slowly than WT cells, and reached the G2 phase with a delay of approximately four hours (Fig. 4C). However, following the induction of DNA damage, CSNAP-depleted cells, unlike the WT cells that displayed a slight delay in progression, remained stalled in the S and G2 phases. This scenario could be due to impaired checkpoint control, rather than exclusively due to a faulty DNA repair mechanism. We therefore validated that the activation of the UV-induced kinase, Chk1, is not dependent on CSNAP, (Figure S6). Similarly, comparison of the colony-forming potential of WT and ΔCSNAP cells following UV irradiation, indicated a significant, 2.7-fold reduction in the number of colonies of cells lacking CSNAP (Fig. 4D). This finding suggests that the accumulation of damaged DNA compromises cell cycle progression and reproductive ability in ΔCSNAP cells.

Next, we determined whether the absence of CSNAP affects DNA damage-induced cellular apoptosis. To this end, we measured the populations of live, early apoptotic and late apoptotic cells in UV-exposed WT and ΔCSNAP cultures four hours post-damage, using flow cytometry. We found that the population of early apoptotic cells following UV exposure is significantly enlarged in WT cells (Fig. 4E), a phenomenon that does not occur at that time point in ΔCSNAP cells, suggesting that the latter fail to efficiently activate the early apoptotic response as WT cells.

Cleavage of PARP1 by caspases is considered to be a hallmark of apoptosis (Chaitanya, Steven et al., 2010, Kaufmann, Desnoyers et al., 1993, Soldani & Scovassi, 2002), and in agreement with the above results, a cleavage product of the protein was detected only in WT but not in ΔCSNAP cells (Fig. 3C). To further examine this phenomenon, we monitored the appearance of the 89 kDa cleavage product of PARP1 following UV irradiation, in a time-dependent manner. We found that in WT cells, the presence of the 89 kDa fragment could already be detected 1 hour following DNA damage (Fig. 4F, Fig. S7). The levels of the cleavage product increased over time, concomitantly with the reduction of the full-length PARP1 (113 kDa) protein. In ΔCSNAP cells, however, the relative abundance of PARP1 was lower than in WT cells, even prior to UV irradiation, and the formation of the 89 kDa cleavage product was only detected after 6.5 hours. Therefore, delayed PARP1 cleavage in ΔCSNAP cells may explain the inability of these cells to activate the early apoptotic response.

Previous studies have shown that CSN is physically recruited to DNA damage sites on the chromatin, and on its path partners with CUL4A^DDB2^ (Fuzesi-Levi et al., 2014, Groisman, Polanowska et al., 2003, Meir et al., 2015). The CSN/ CUL4A^DDB2^ association is rapidly relieved at the DNA lesion site, to induce activation of the CUL4A^DDB2^ complex. Thus, we examined the associations of both CSN^ΔCSNAP^ and the WT complex with CUL4A^DDB2^ components, following the induction of DNA damage. Time-course analysis of DDB2 pull-downs from chromatin-bound fractions following UV irradiation indicated that, as expected, both DDB2 and DDB1 are rapidly recruited to chromatin (Fig. 4G). In WT cells, we could detect CSN release from the DDB2 complex following UV irradiation. However, in ΔCSNAP cells, although release was observed after 20 minutes it was less significant and rapid restoration of CSN/DDB2 interaction was detected, compromising the activation of the DNA damage response through CUL4A^DDB2^. This is observation is consistent with our in vitro binding data, which showed a stronger binding between Cul1/Rbx1 and CSN^ΔCSNAP^ versus CSN (Fig. 1E) and between CSN3 and DDB2 (Fig. 1B and C). Overall, these results support the view that the affinity of CSN for CRL complexes is enhanced, in the absence of CSNAP.

## Discussion

Here, we investigated the functional contribution of CSNAP, the smallest and last to be discovered CSN subunit to the steric and catalytic functions of the CSN. We find that CSNAP attenuates CSN binding interactions with CRL (Fig. 5). Efficient dissociation from CRL assemblies is essential for reconfiguration of new CRL compositions in order to respond to changing regulatory inputs. Therefore, a hypothesis emerging from this study is that the increased affinity of CSN^ΔCSNAP^ for CRLs will affect the dynamic plasticity of CRL configuration. Indeed, we find that the absence of CSNAP alters cell cycle progression and reduces cellular viability. In addition, the impaired DNA damage response following UV irradiation of CSN^ΔCSNAP^ indicates a reduced capacity of ΔCSNAP cells to adapt to cellular stimuli. Together these results show that CSNAP contributes to the steric regulation of CRL by CSN, with global cellular effects.

**Figure 5.**
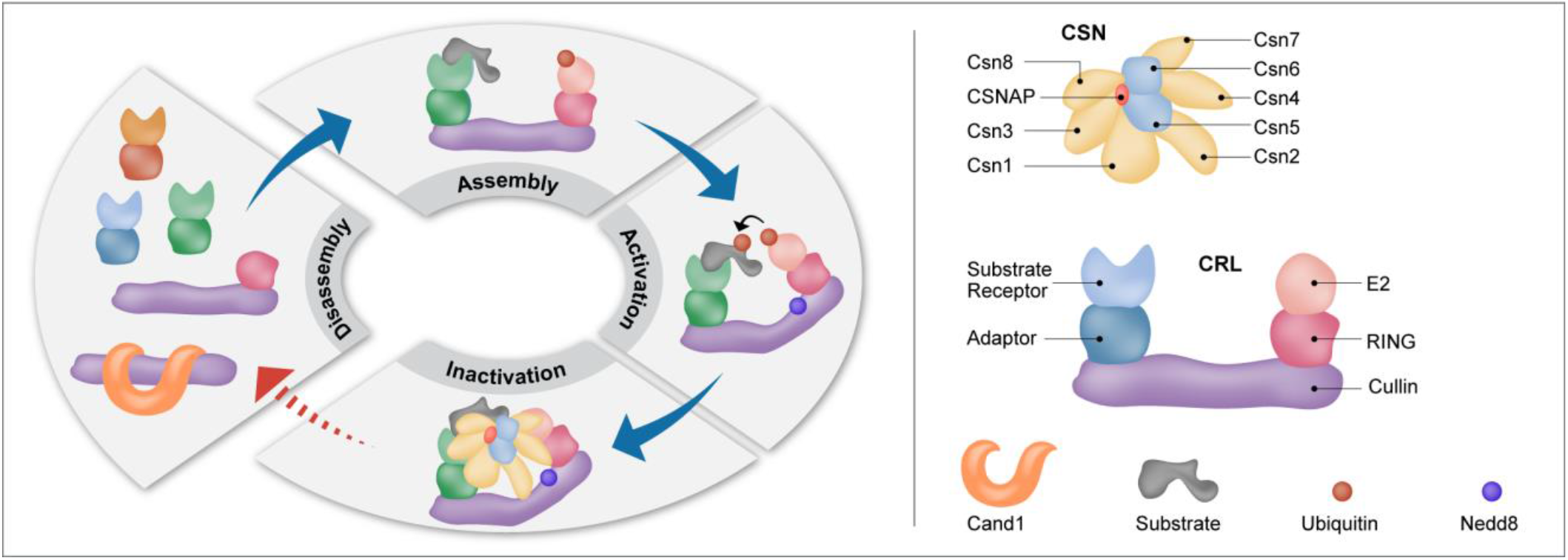
CSNAP influences the strength of the CSN-CRL interaction. Diagram representing the CRL cycle. CRLs form dynamic complexes with various adaptors and substrate receptors. The conjunction of Nedd8 to a conserved lysine residue in the cullin subunit, induces a conformational change that activates the CRL complex, promoting ubiquitin transfer to the substrate. The CSN complex inactivates CRL assemblies by two independent mechanisms, catalytic and non-catalytic. The first involves catalytic removal of the Nedd8 conjugate, while the second is mediated through physical binding to CRLs, sterically precluding interactions with E2 enzymes and ubiquitination substrates. Subsequently, after CSN dissociation, CRLs can be disassembled and assembled into new configurations, or bind Cand1. This cycle enables CRL adaptation according to cellular need, enabling specific substrates to be ubiquitinated. Our results indicate that CSNAP reduces the affinity of CSN for CRL, thus enabling efficient disassembly and remodeling of CRL complexes. In the absence of CSNAP, the disassembly and assembly steps of the cycle are compromised, as designated by the red dashed lines, affecting the reconfiguration of CRL assemblies, and their ability to respond to cellular stimuli.

Our data indicate that the *K_d_* for CSN binding to Cul1 is at least 3 fold higher than for the complex lacking CSNAP (Fig. 1E). Given that the *K_d_* is in the micromolar range, and that the cellular cullin and CSN concentrations are ~2.2 and 0.45 μM, respectively (Mosadeghi et al., 2016), the change in *K_d_* would be expected to impact the free CSN and CRL pools. Unneddylated cullins bind Cand1 or Cand2, the F-box protein exchange factors that mediates CRL recycling (Liu et al., 2018, Pierce, Lee et al., 2013, Schmidt, McQuary et al., 2009) (Fig. 5). A portion of unneddylated cullins, however, were shown to remain unbound to Cand1 (Bennett, Rush et al., 2010, Liu, Zhou et al., 2017, Schmidt et al., 2009) and some SCF substrates are efficiently degraded independently of these exchange factors (Liu et al., 2018, Scott & Schulman, 2018). Hence, free unneddylated cullins may be directly available for configuration of new CRL modules.

In line with this assumption, it was demonstrated recently that the presence of unneddylated Cul1 is important for maintaining the substrate receptor pool and promoting rapid assembly and activation of Cull-Skpl-F-box complexes (Liu et al., 2017). Moreover, prolonging CSN-CRL interaction using irreversible neddylation inhibits CRL activity (Scherer, Ding et al., 2016). Likewise, strengthening the CRL-CSN interaction using the metabolite inositol hexakisphosphate promotes CRL inactivation (Scherer et al., 2016). Taken together with the present results, it is reasonable to conclude that modulation of CSN-CRL binding is an important mode of CRL regulation.

The recent crystal structure of free CSN^ΔCSNAP^ indicated that PCI and MPN subunits form largely distinct substructures (Lingaraju, Bunker et al., 2014). The six PCI subunits comprise the base of the CSN, with their C-terminal ends forming an elaborate bundle above which the heterodimer CSN5/CSN6 sits. Previously, we discovered that CSNAP tethers together these two distinct structural elements, by mutually binding CSN5/CSN6, and the PCI subunit CSN3. Both CSN5 and CSN3 directly interact with the CRL assembly, CSN3 with the substrate receptor (Cavadini, Fischer et al., 2016, Enchev et al., 2012), and CSN5 with the Nedd8 cullin modification (Cope et al., 2002). Thus, it is likely that through these interactions, CSNAP modulates CSN-CRL interactions. Unraveling the precise structural contribution of CSNAP to CSN-CRL binding affinity awaits high-resolution structural analyses; however, it is reasonable to speculate that CSNAP shifts the CSN conformational equilibrium toward low affinity states.

Given that CRLs are involved in regulating numerous cellular processes, including cell-division cycle and cellular proliferation, the correlation between aberrant CRL function and cancer is not surprising, making this system an attractive target for therapeutic intervention (reviewed in (Kitagawa & Kitagawa, 2016, Wang, Liu et al., 2014, Zhao & Sun, 2013)). For example Cul4/CRBN has been implicated as the target of the anti-myeloma agent lenalidomide (Lu, Middleton et al., 2014), and the neddylation inhibitor MLN4924 is an anti-cancer drug currently in clinical trials (Soucy et al., 2009). As a direct regulator of CRLs, the CSN constitutes another objective for drug development, with a major focus on inhibiting CSN5, the catalytic subunit (Altmann, Erbel et al., 2017, Cope & Deshaies, 2006, Lauinger, Li et al., 2017, Lee, Judge et al., 2011, Pulvino, Chen et al., 2015, Schlierf, Altmann et al., 2016). Our latest findings lead us to propose CSNAP as a new therapeutic avenue. Preventing CSNAP integration within the CSN complex would be expected to impair cell cycle progression and the adaptive response to oncogenic stress conditions.

## Materials and Methods

### Cell cultures, transfections, and UV-C exposure

HAP1 WT and ΔCSNAP CRISPR cell lines were purchased from Haplogene GmbH, Austria, and cultured in a humidified CO_2_ incubator at 37 °C in Iscove’s Modified Dulbecco’s Medium (IMDM) supplemented with 10% fetal calf serum, penicillin-streptomycin and Mycozap (Lonza). HAP1 cells were transfected with Hyg-CSNAP-Cerulean, Hyg-ΔN-CSNAP-Cerulean, or Hyg-FBXL15-FLAG, using the JetPrime reagent (Polyplus). Cerulean-expressing cell lines were isolated and sorted for low-medium expression levels by fluorescence-activated cell sorting (FACSAria Fusion; BD Biosciences), and expanded in complete IMDM. For UV treatments, plates were washed twice with PBS, and after removal of the liquid, were illuminated with 5 or 20J/m^2^ UV-C light.

### Immunoprecipitation and FLAG-pull down

For immunoprecipitation experiments, HAP1 cells were lysed in 50 mM Tris pH 7.4, 150 mM NaCl, 0.5% NP40, phosphatase inhibitors (5 mM Na-o-vanadate, 4 mM Na-pyrophosphate and β-glycerophosphate) and protease inhibitors inhibitors (1 mM PMSF, 1 mM benzamidine, 1.4 μg/ml pepstatin A. 0.25–1 mg total protein was incubated with 10μl anti-CSN3 (Abcam ab79398), anti-DDB2 (Santa Cruz sc-81246) or 35μ1 anti-FLAG resin (Sigma A2220) overnight. For immunoprecipitation of the CSN3 or DDB2 antibody 35 μl protein G sepharose slurry was added for 1 hour. Bound proteins were washed and eluted with 2x Laemmli sample buffer. Chromatin-bound proteins were purified as previously described (Fuzesi-Levi et al., 2014), using 50 μg/ml digitonin instead of NP40 in the hypotonic lysis buffer. 250 μg of chromatin bound fraction was suspended in 250 μl TBS and rotated overnight at 4 °C with 5 μl of anti-DDB2 (Santa Cruz sc-81246). Then 30 μl of TBS equilibrated Protein G Sepharose resin (GE) was added for 1 hour, and after 3 washes of 300 μl TBS bound proteins were eluted in 35 μl 2x Laemmli sample buffer.

### Fluorescence assays

Purification of recombinant CSN^ΔCSNAP^ complexes was performed as described in (Enchev et al., 2012). The production of CSN and CSN^5H138A^ involved the generation of a pFBDM vector containing CSN1/His6-CSN5/CSN2/StrepII2x-CSN3/CSNAP or CSN1/His6-CSN5^H138A^/CSN2/StrepII2x-CSN3/CSNAP respectively. Baculoviruses produced from each of these vectors were used to co-infect HighFive insect cells with a baculovirus expressing CSN4/CSN7b/CSN6/CSN8 to produce the full complexes. The fluorescent assays to determine the affinity of the CSN complexes for Cul1-dansyl/Rbx1 variants and their deneddylation activity were performed as described in (Mosadeghi et al., 2016).

### Western blots

Proteins were separated on 12% SDS-PAGE, and transferred to PVDF membranes. Primary antibodies used for detection: anti-CSN1 (Enzo PW8285), anti-CSN2 (Abcam ab10462), anti-CSN3 (ab79398), anti-CSN5 (ab495 and ab118841), anti-CSN6 (PW 8295), anti-CSN8 (BML-PW8290), anti-PDCD4 (ab80590), anti-cullin1 (ab75817), anti-cullin2 (ab166917), anti-cullin3 (ab75851), anti-cullin4AB (ab76470), anti-cullin5 (ab184177), anti-DDB1 (Bethyl A300–462A), anti-DDB2 (ab181136), anti-PARP1 (sc-8007), anti-FLAG (Sigma F3165), anti-pChk1 (Cell signaling 2341), anti-GAPDH (Millipore MAB374), anti-vimentin (ab92547), anti-NQO1 (ab28947), anti-tubulin (ab184613), and anti-histone 3 (ab24834). All Western blot analyzes were repeated at least three times.

### Comet assay

Alkaline single-cell electrophoresis was performed, according to the protocol from Trevigen. HAP1 cells were treated with 20J/m^2^ UV-C light, and 6 hours post-exposure cells were trypsinized, counted, and suspended in ice-cold PBS (-Ca/-Mg) to a density of 2×10^5^ cells/ml. Fifty μl of cell suspension were mixed with 450 μl LM-agarose (Trevigen), and 50 μl of the mix was pipetted onto the comet slide, and incubated in the dark at 4 °C for 30 minutes. Slides were immersed in a lysis solution (Trevigen) for 1 hour at 4 °C, and then equilibrated to an alkaline electrophoresis solution (300 mM NaOH, 2 mM EDTA, pH>13) for 20 minutes at room temperature. Slides were run at 1 V/cm (~300 mA constant) in ice-cold alkaline electrophoresis solution for 30 minutes and then neutralized for 5 minutes in 400 mM Tris, pH7.5, rinsed in distilled water, immersed in 70% ethanol for 5 minutes and dried at room temperature. DNA was stained with SYBR Gold (Invitrogen); the slides were then dried completely, prior to imaging. Images were acquired using an inverted Nikon microscope (Eclipse Ti, Nikon, Japan) using a 20x objective, and with a cooled electron-multiplying charge-coupled device camera (iXon Ultra, Andor, Ireland). Comet parameters were analyzed using the CASP comet software. At least 74 cells were analyzed per sample, in 3 biological samples, in duplicates.

### Colony-forming assay

Untreated or 20J/m^2^ UV-C exposed WT and ΔCSNAP cells were trypsinized, counted, and plated in 10 cm tissue culture dishes in triplicates. For untreated cells, 100 cells, and for UV-illuminated cells, 5,000 cells were plated per dish. All experiments were done in triplicates. Dishes were incubated in normal growth conditions for 8 days. The plates were then washed twice with PBS, dried, and stained with 0.15% Crystal violet in methanol for 3 minutes, rinsed with tap water, and air-dried before scanning. Colony counts were measured using OpenCFU software.

### Cell cycle analysis

Cells were synchronized to G1/S phase using double thymidine block as previously described (Fuzesi-Levi et al., 2014). UV treated cells were exposed to 5 J/m^2^ UV-C at release, and fixed with ethanol at different time points. Cell cycle phases were assessed by flow cytometry (LSRII, BD Biosciences) following propidium iodide staining. Asynchronous cells were analyzed using propidium iodide and BrdU double staining. 10^5^ cells were denatured after fixation using 2N HCl, 0.5% Triton X-100 in PBS for 30 minutes, neutralized in 0.1 M Na_2_B_4_O_7_ pH 8.5 and incubated with 5 μl of anti-BrdU-FITC (eBioscience 11–5071–41) in 1% BSA, 0.5% Triton X-100 in PBS for 1 hour. Cells were washed in 1% BSA in PBS, resuspended in PBS containing 50 μg/ml propidium iodide and 50 μg/ml RNase A, and analysed in FACSAria Fusion flow cytometer (BD Biosciences).

### Measurement of viable and dead cell populations

Determination of percentage of live, early apoptotic and late apoptotic/necrotic cells in WT and ΔCSNAP cultures was performed using Annexin V-FITC Apoptosis Detection Kit (APOAF-20TST, Sigma) by flow cytometry. Single cells were analyzed for AnnexinV-FITC and propidium iodide fluorescence. Early apoptotic cells (annexin V positive, PI negative), late apoptotic/necrotic/dead cells (annexin V positive, PI positive), and live/viable cells (annexin V negative, PI negative) were gated and quantified.

### Resazurin assay

WT, ΔCSNAP, ΔCSNAP-Cerulean and ΔCSNAP-AC-Cerulean expressing cells were trypsinized, counted, and seeded in 4 replicates at a cell density of 5000 cells/well in 24 well plates. Cells in one plate were seeded directly to 30 μ,g/ml resazurin containing growth medium, and fluourescence intensity (540/600nm) was measured after 2 hours, for initial proliferation value. The other plates were either UV-exposed at 20 J/m^2^ 24 hours after seeding or left untreated. Two or 4 hours post-UV the growth medium was changed to 30 μg/ml resazurin containing medium, incubated for 2 hours, and fluorescence was measured. Proliferation was calculated at each time point normalizing to the initial proliferation value.

### SILAC

HAP1 cells were grown in SILAC IMDM (Invitrogen) with 10% dialyzed fetal calf serum (Biological Industries, 04–011–1A) supplemented with 2 5 μg/ml light L-lysine and L-arginine (Sigma) or 25 μg/ml heavy L-lysine (L-Lys8-CNLM-291-H-1, Cambridge Isotopes) and L-arginine (L-Arg10-CNLM-539-H-1, Cambridge Isotopes) each, and labeling was swapped between WT and ΔCSNAP cells. Cells were incubated with 5μM MG132 for 4 hours before harvesting. Samples were prepared, as previously described (Udeshi, Svinkina et al., 2013). Briefly, the samples were lysed using 8 M urea, mixed at a 1:1 protein:protein ratio, and digested with trypsin, followed by a desalting step. The resulting peptides were fractionated offline using high pH-reversed phase chromatography, followed by enrichment for K-ε-GlyGly using the Cell Signaling PTMScan^®^ Ubiquitin Remnant Motif (K-ε-GG) Kit #5562 (antibody-based). Each fraction was then analyzed, using online nanoflow liquid chromatography (nanoAcquity) coupled to high-resolution, high-mass accuracy mass spectrometry (Fusion Lumos). Raw data was processed with MaxQuant v1.5.5.1. The data was searched with the Andromeda search engine against the human proteome database appended with common lab protein contaminants, and allowing for GG modifications of lysines. The ratio of H/L (heavy to light) ratio was calculated, and results were log-transformed. The datasets of four (two label swap) experiments were combined in the way, that the experiment with the largest number of proteins identified was merged with proteins that had data for the same proteins and modification sites from the three other experiments. Genes corresponding to proteins that showed fold change above 1.5 or below 0.66 in each of the four experiments were filtered, and were selected only if appeared at least two out of the four experiments (from non-unique proteins only the first in the gene list was included). The resulting protein/gene list from was analyzed using EnrichR/Reactome 2016 or Webgestalt/Reactome 2016 for overrepresentation, and results were filtered using a cut off of adjusted p value or FDR < 0.05.

### Label-free quantitation

WT and ΔCSNAP cells were lysed in 50 mM Tris, pH 7.4, 150 mM NaCl, 0.5% NP40, supplemented with phosphatase and protease inhibitors as described above. One mg total protein was used for immunoprecipitation, using anti-CSN3 antibody (ab79698) as described above in 3 biological replicates. Proteins were eluted by 75 μl of 0.1 M glycine-HCl, pH 2.5. The beads were washed in 25 mM Tris pH7.4, 150 mM NaCl (TBS), and subjected to on-bead tryptic digestion as follows: 8 M urea in 0.1 M Tris, pH 7.9, was added onto TBS washed beads, and incubated for 15 minutes at room temperature. Proteins were reduced by incubation with dithiothreitol (5 mM; Sigma) for 60 minutes at room temperature, and alkylated with 10 mM iodoacetamide (Sigma) in the dark for 30 minutes at room temperature. Urea was diluted to 2M with 50mM ammonium bicarbonate. Trypsin (250 ng; Promega; Madison, WI, USA) was added and incubated overnight at 37 °C, followed by addition of 100 ng trypsin for 4 hour at 37 °C. Digestions were stopped by addition of trifluoroacetic acid (1% final concentration). Following digestion, peptides were desalted using Oasis HLB μElution format (Waters, Milford, MA, USA), vacuum-dried, and stored at −80°C until further analysis.

ULC/MS grade solvents were used for all chromatographic steps. Each sample was loaded using split-less nano-Ultra Performance Liquid Chromatography (10 kpsi nanoAcquity; Waters). The mobile phase was: A) H_2_O + 0.1% formic acid and B) acetonitrile + 0.1% formic acid. Sample desalting was performed online, using a reversed-phase Symmetry C18 trapping column (180 μm internal diameter, 20 mm length, 5 μm particle size; Waters). The peptides were then separated using a T3 HSS nano-column (75 μm internal diameter, 250 mm length, 1.8 μm particle size; Waters) at 0.35 μL/min. Peptides were eluted from the column into the mass spectrometer, using the following gradient: 4% to 30% B for 55 min, 30% to 90% B for 5 min, maintained at 90% for 5 min, and then back to initial conditions. The nanoUPLC was coupled online through a nanoESI emitter (10 μm tip; New Objective; Woburn, MA, USA) to a quadrupole orbitrap mass spectrometer (Q Exactive Plus, Thermo Scientific), using a FlexIon nanospray apparatus (Proxeon).

Data was acquired in data-dependent acquisition (DDA) mode, using a Top20 method. MS1 resolution was set to 70,000 (at 400 m/z), mass range of 300–1650 m/z, AGC of 3e6, and maximum injection time was set to 20 msec. MS2 resolution was set to 17,500, quadrupole isolation 1.7 m/z, AGC of 1e6, dynamic exclusion of 30 sec, and maximum injection time of 60 msec. Raw data was imported into Expressionist^®^ software version 9.1.3 (Genedata), and processed as described here. The software was used for retention time alignment and peak detection of precursor peptides. A master peak list was generated from all MS/MS events, and sent for database searching using Mascot v2.5.1 (Matrix Sciences). Data was searched against the human sequences UniprotKB (http://www.uniprot.org/), appended with the CSNAP sequence and common laboratory contaminant proteins. Fixed modification was set to carbamidomethylation of cysteines, and variable modifications were set to oxidation of methionines and deamidation of N or Q. Search results were then filtered using the PeptideProphet algorithm, to achieve a maximum false discovery rate of 1% at the protein level. Peptide identifications were imported back to Expressionist to annotate identified peaks. Quantification of proteins from the peptide data was performed, using an in-house script. Data was normalized, based on the total ion current. Protein abundance was obtained by summing the three most intense, unique peptides per protein. A Student’s t-test, after logarithmic transformation, was used to identify significant differences (>1.5-fold) across the biological replica. Fold changes were calculated based on the ratio of arithmetic means of the case versus control samples.

### Total proteome analysis and bioinformatics

WT and ΔCSNAP cells were lysed in SDT buffer (4% SDS, 100 mM Tris/HCl, pH 7.6, and 0.1 M dithiothreitol) and subjected to tryptic digestion, using a FASP™ Protein Digestion Kit (Expedeon). The resulting peptides were desalted and analyzed on the LC-MS instrument (Q-Exactive Plus) in DDA mode. The raw data was processed in Expressionist by Genedata, using Mascot as the search engine against the uniprot human proteome database, and common protein contaminants. Identifications were filtered to a maximum of 1% FDR on both the peptide and protein levels. Protein inference was performed by an in-house script. Overall, about 4,000 proteins were identified and quantified. Proteomics data, after logarithmic transformation and flooring, were analyzed by two-way ANOVA using two factors, strain and UV treatment, as well as their interaction. Proteins with a p value of <0.05 and an absolute fold change >1.5 were considered to be differentially expressed. The proteins were filtered to keep those that had an absolute fold change of at least 1.5 and a p-value of <0.05 in at least one of the following pairwise comparisons: 1. WT UV/WT untreated; 2. ΔCSNAP UV/ ΔCSNAP untreated; 3. ΔCSNAP untreated/ WT untreated; 4. ΔCSNAP UV/ WT UV. The log intensities of the 347 proteins that passed these criteria (according to ANOVA analysis with all samples, after flooring was used). Intensities were clustered using the k-means algorithm, with Pearson dissimilarity as the distance measure to 5 clusters. log_2_ intensities were standardized, so that each protein displayed zero mean and unit standard deviation. The proteins in each cluster could be obtained by filtering the Excel file. Enrichment analysis of the filtered protein list was performed using Webgestalt overrepresentation analysis using Reactome 2016 pathway as functional database against the protein coding database as reference set, and results were filtered for adjusted p value <0.05.

All raw data, peak lists and identifications were deposited to the ProteomeXchange Consortium http://proteomecentral.proteomexchange.org) via the PRIDE partner repository.

### Native mass spectrometry

Nano-electrospray ionization mass spectrometry (MS) experiments were performed on a modified Q Exactive Plus Orbitrap EMR (Ben-Nissan, Belov et al., 2017). Prior to MS analysis, 25 μl of the samples (~ 16 μM) were buffer-exchanged into 0.5 M ammonium acetate (pH 7), using Bio-Rad Biospin columns. Two μl of the buffer-exchanged samples were mixed with 1 μl MeOH 40%, to reach a final concentration of 13%. Proteins were loaded into gold coated nano-ESI capillaries, prepared in house from borosilicate glass tubes (Kirshenbaum, Michaelevski et al., 2010). For the reconstitution of the CSN complex, after buffer exchange, 2 μl of CSN^ΔCSNAP^, were incubated for 3 h on ice with 2 μl of a synthetic CSNAP peptide, dissolved to 50 μM in 250 mM ammonium acetate. The conditions within the mass spectrometer were adjusted to optimize signals of the intact CSN and preserve non-covalent interactions. The instrument was operated in positive mode at capillary voltage of 1.7 kV. Argon was used as the collision gas in the higher energy collision-induced dissociation (HCD) cell. Resolution was set to 8750. Forevacuum was set to 1.5 mbar and the trapping gas was set to 3, corresponding to pressures of 8.8×10–5 and 1.7×10–10 mbar in the HV and UHV regions, respectively. Flatapole bias was set to transmission at 1.5 V. Bent flatapole and axial gradient were set to DC 2.2 V and 37.2 V, respectively. HCD cell bias was set to 150 V. Spectra are shown with no smoothing and without background subtraction.

## Acknowledgments

We thank Dieter A. Wolf for comments on the manuscript, and Dr. Ron Rotkopf for the statistical analysis. M.S. is grateful for the financial support of the US National Institutes of Health, grant no. GM121834. M.S. is the incumbent of the Aharon and Ephraim Katzir Memorial Professorial Chair. G.F. is the Incumbent of the David and Stacey Cynamon Research fellow Chair in Genetics and Personalized Medicine

## Author contribution

M.G.F.-L., G.B-N., R.I.E., M.P. and M.S. designed the experiments and analyzed the data. M.G.F.-L. performed the cell biology and biochemistry experiments. M.G.F.-L and T.M.S performed the flow cytometry, and M.G.F.-L., and R.N. performed the microscopy experiments. G.B-N. performed the mass spectrometry and R.I.E. the *K_d_* measurement experiments. Y.L. and M.K. performed the SILAC and label-free proteomics analysis. G.F. performed the bioinformatics analysis of the proteomics data. M.G.F.-L, G.B.-N., and M.S. wrote the manuscript.

## Supplemental Information

## Supplemental Figures

**Figure S1.**
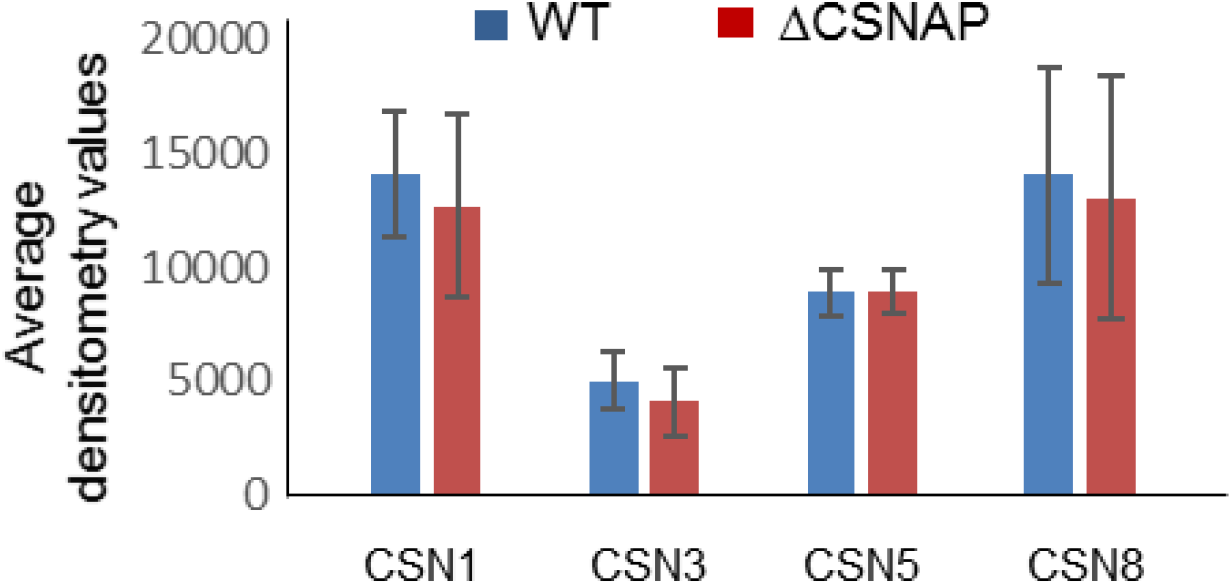
The levels of CSN subunits are comparable in WT and ΔCSNAP cells. Quantification of CSN subunit levels from three independent experiments with standard errors. (see Fig.1D).

**Figure S2.**
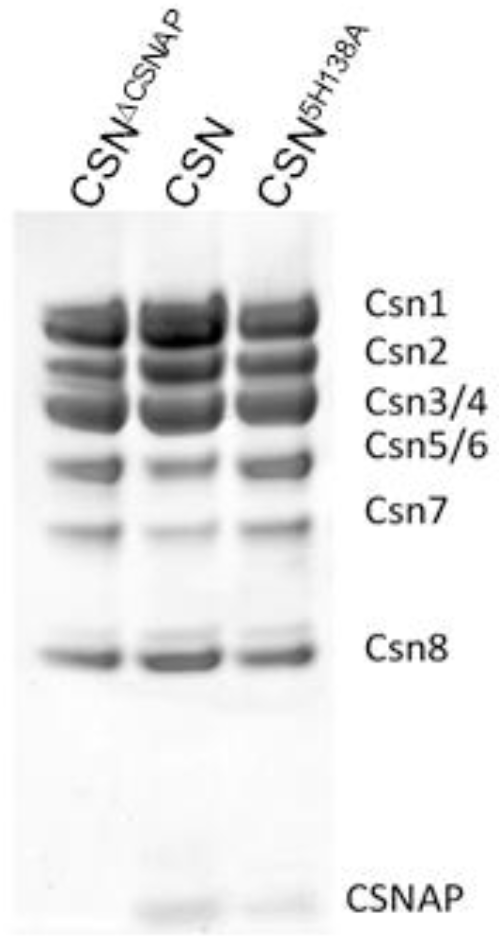
Recombinant CSN complexes. Purified WT and mutant (CSN5^H138A^) CSN complexes expressed with or without CSNAP in insect cells.

**Figure S3.**
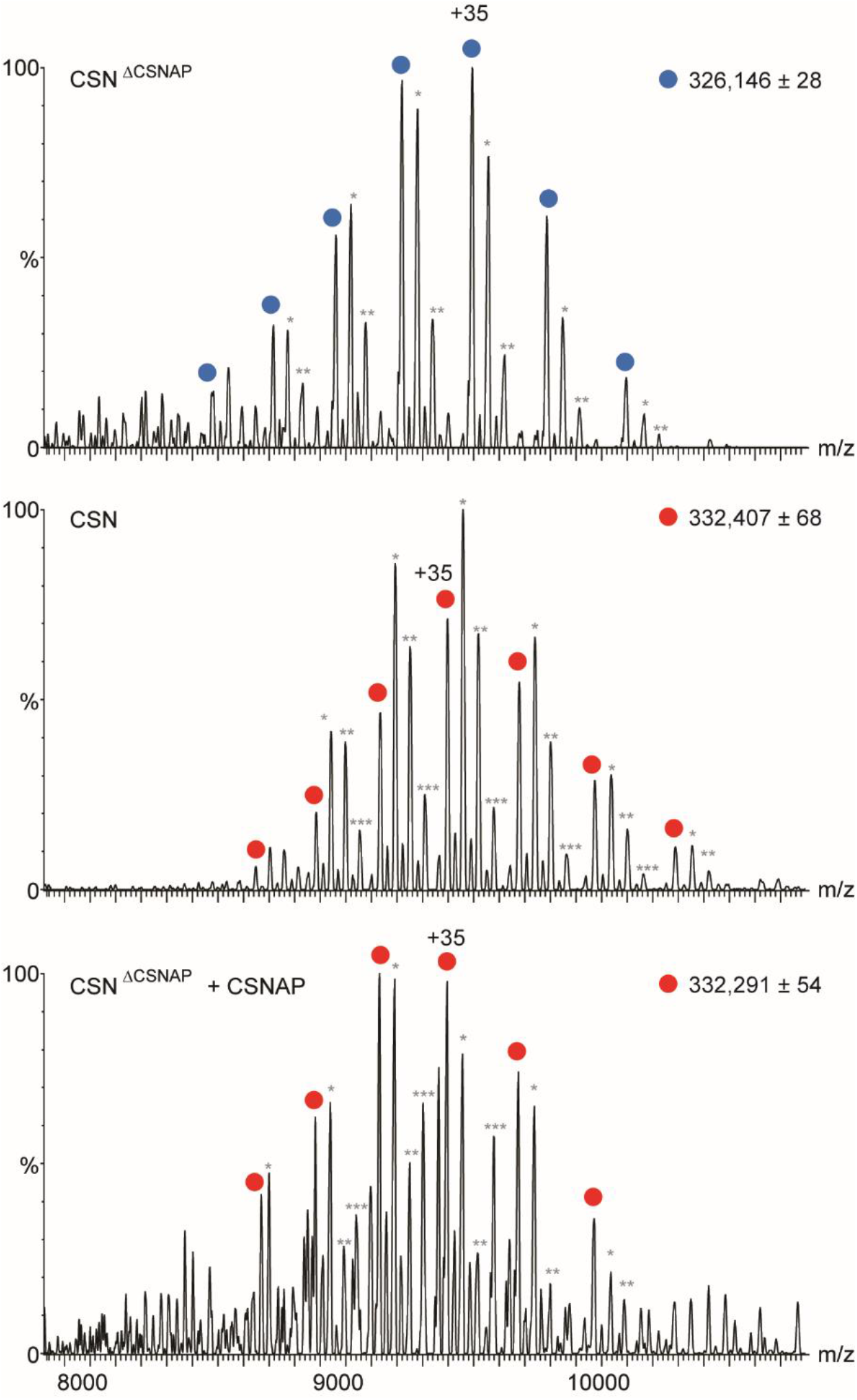
Recombinant CSN and CSN^ΔCSNAP^ complexes form intact complexes. Native mass spectrometry analysis of recombinant CSN complexes, purified from insect cells. The mass difference between cSN^ΔCSNAP^ (top panel, blue circles) and CSN (middle panel, red circles) coincides nicely with the mass of CSNAP, indicating that the intact complex contains a single and stoichiometric CSNAP subunit. The bottom panel shows that when the CSN^ΔCSNAP^ is supplemented with a synthetic CSNAP peptide, it is reconstituted into the intact, CSN complex (red circles). Asterisks indicate populations with purification adducts.

**Figure S4.**
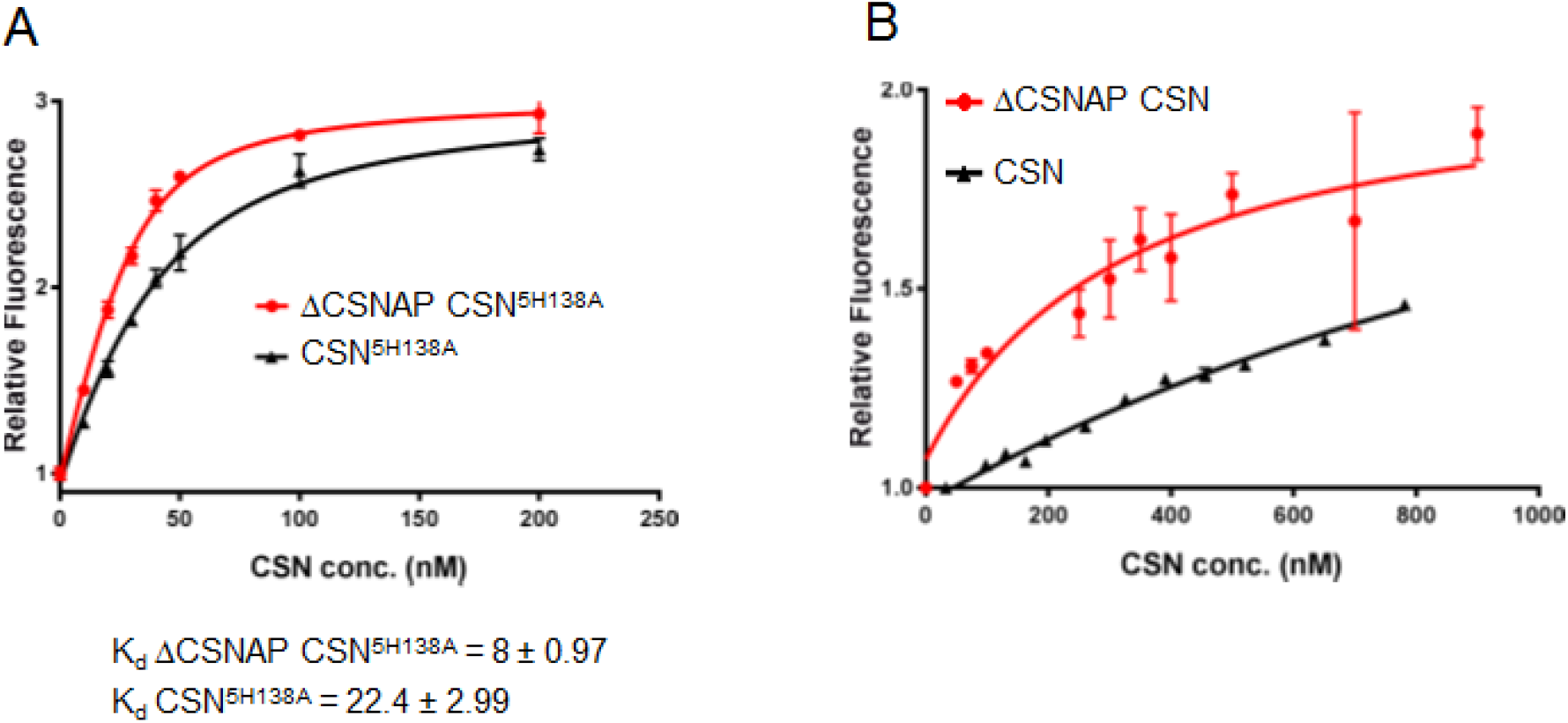
The absence of CSNAP increases the affinity towards Cul1. *K_d_* for CSN and CSN^ΔCSNAP^ complexes were determined by measuring the change in dansyl fluorescence using the well characterized H138A CSN5 mutation (CSN^5H138A^) (A), the WT CSN5 subunit (B), and the dansyl-labeled Cul1-N8/Rbx1CSN complexes. All binding measurements were carried out in triplicate and error bars represent standard deviation.

**Figure S5.**
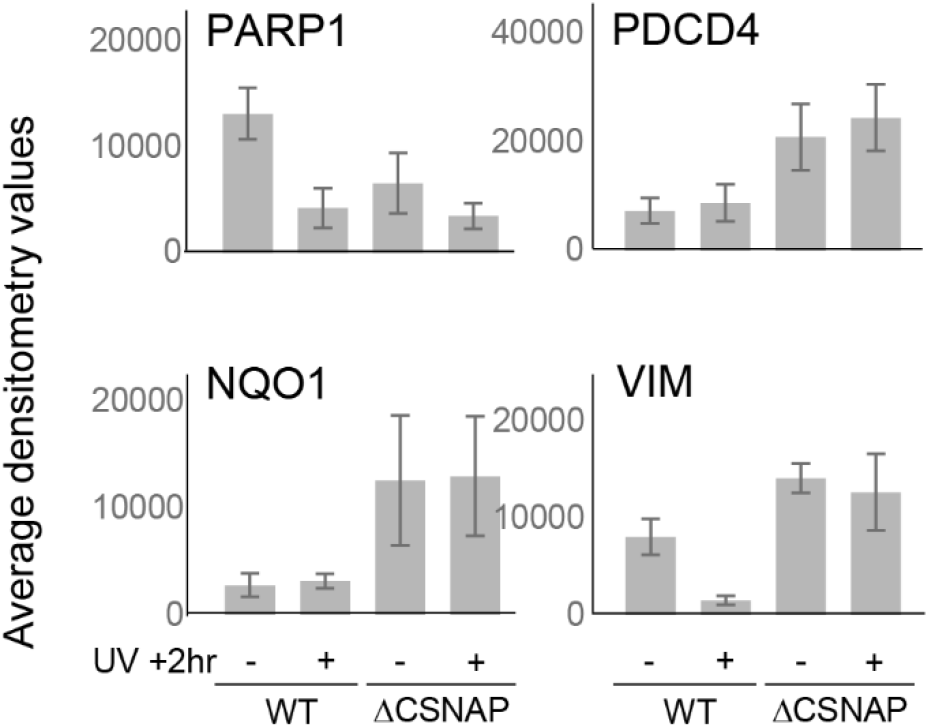
Differences in protein expression levels among WT and ΔCSNAP cells. Densitometry analysis of PARP1, PDCD4, NQO1 and vimentin (shown in Fig.3C). Values are average of three independent experiments; error bars represent standard errors.

**Figure S6.**
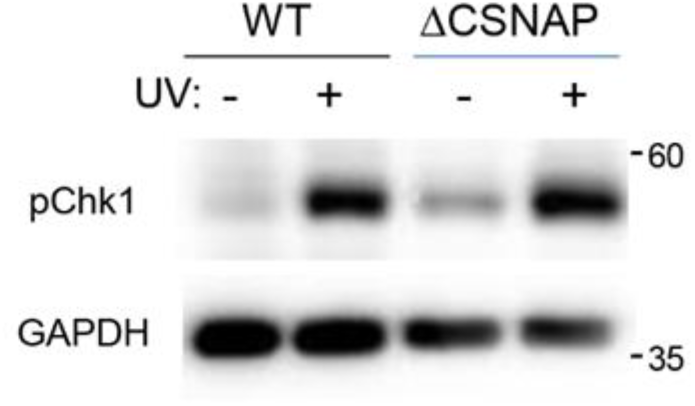
Checkpoint control is unaffected in cells lacking CSNAP. Untreated and UV-exposed WT and ΔCSNAP cells were lysed four hours post-damage, and phosphorylation of Chk1 (Ser345) was compared. In response to UV irradiation, Chk1 is phosphorylated, as expected. Thus, phosphorylation of Chk1 is not affected in cells lacking CSNAP.

**Figure S7.**
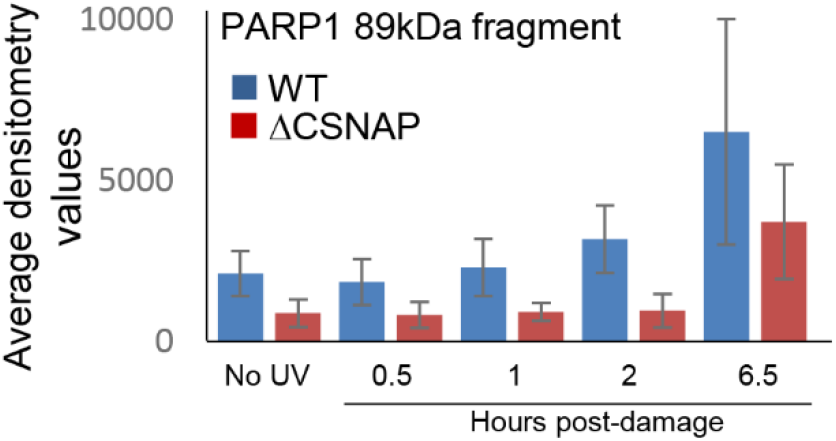
PARP1 cleavage is delayed in ΔCSNAP cells. Densitometry analysis of the full length PARP1 and its 89 kDa cleavage fragment (shown in Fig.4F). Values are average of three independent experiments; error bars represent standard errors.

**Table S1.**
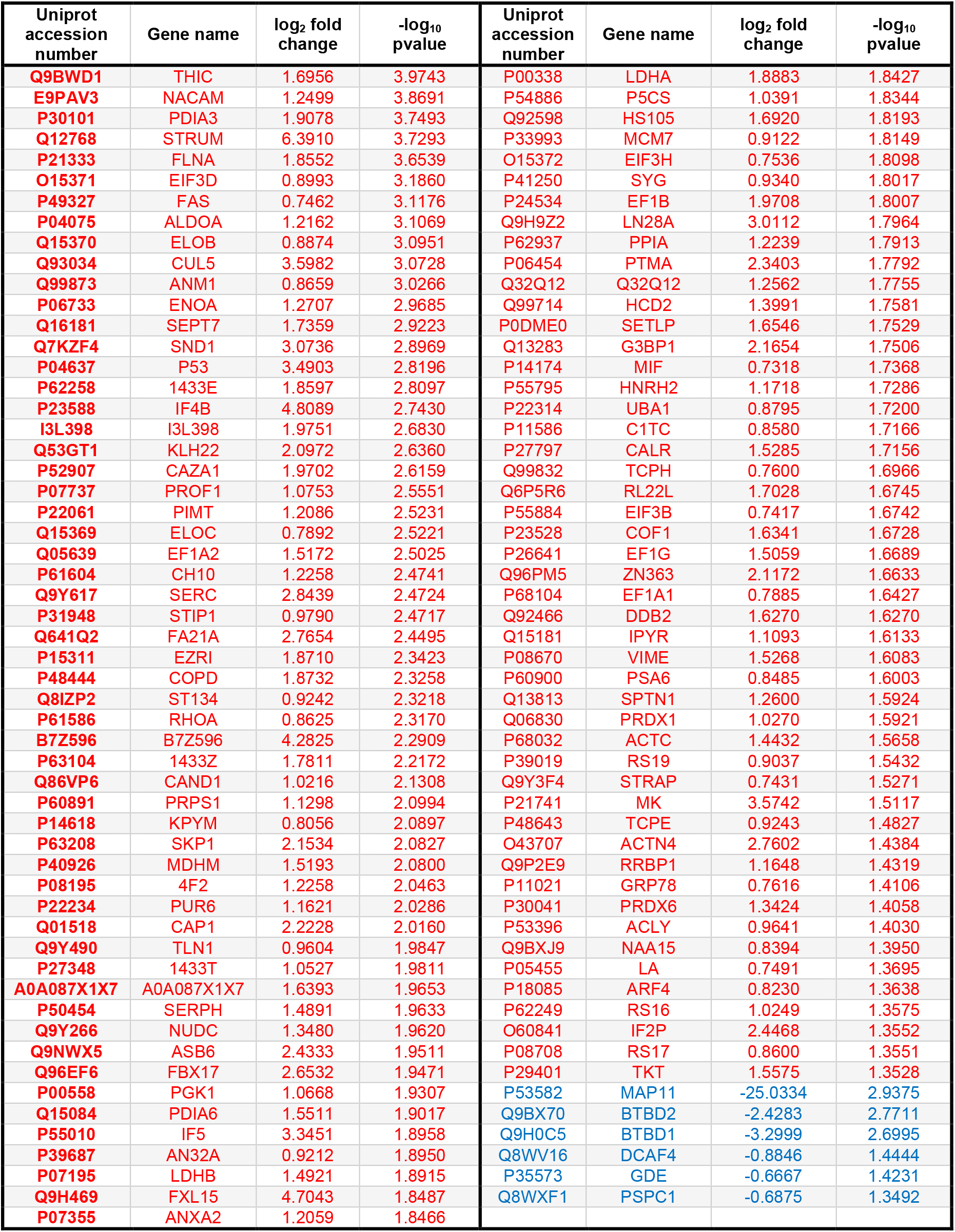
Proteins significantly enriched in ΔCSNAP cells in comparison to WT cells, yielded by label-free proteomic analysis following immunoprecipitation through CSN3. (-log_10_ p value>1.3; log_2_ fold changes x < −0.58 (blue) and x > 0.58 (red)).

**Table S2A.** The table shows the differentially ubiquitinated proteins obtained in at least two out of four experiments, with log_2_ fold change (ΔCSNAP / WT) above 1.5 in SILAC proteomic analysis.

| Gene Name | Uniprot accession number | Gene Name | Uniprot accession number | Gene Name | Uniprot accession number |
| --- | --- | --- | --- | --- | --- |
| OGT | O15294 | RPS3 | P23396 | SLC1A4 | P43007 |
| KIF1A | Q12756 | NBR1 | Q14596 | MAP1B | P46821 |
| USP9X | Q93008 | HIF1A | Q16665 | VAMP7 | P51809 |
| TOM1 | O60784 | HSD17B12 | Q53GQ0 | SLC12A2 | P55011 |
| RPN1 | P04843 | PI4K2A | Q9BTU6 | YBX1 | P67809 |
| SLC3A2 | P08195 | SMAP | O00193 | SLC7A5 | Q01650 |
| VIM | P08670 | BSG | P35613 | TRAF2 | Q12933 |
| PARP1 | P09874 | SLC19A1 | P41440 | TRAF3 | Q13114 |
| XRCC5 | P13010 | SLC6A8 | P48029 | PEG10 | Q86TG7 |
| RBMX | P38159 | ACLY | P53396 | APH1B | Q8WW43 |
| PEX19 | P40855 | SLC16A1 | P53985 | EPS15L1 | Q9UBC2 |
| NEDD4 | P46934 | RAD23A | P54727 | VTI1B | Q9UEU0 |
| UBE2N | P61088 | RAD23B | P61956 | SSR3 | Q9UNL2 |
| RAB1A | P62820 | YWHAZ | P63104 | ATXN10 | Q9UBB4 |
| VAMP2 | Q15836 | MSMO1 | Q15800 | HIST1H1E | P10412 |
| PPP2CA | P62714 | CCDC50 | Q8IVM0 | HIST1H1C | P16403 |
| ZMYM3 | Q14202 | ARL8A | Q96BM9 | ARL8B | Q9NVJ2 |
| PCM1 | Q14202 | ITCH | Q96J02 | UBXN7 | O94888 |
| STX12 | Q15154 | FANCI | Q9NVI1 | FADS2 | O95864 |
| ANKRD13A | Q8IZ07 | NSFL1C | Q9UNZ2 | PCNA | P12004 |
| PDCD6IP | Q8WUM4 | ERH | P84090 | BCAP31 | P51572 |
| SERINC1 | Q9NRX5 | SLC1A5 | Q15758 | RPS27A | P62979 |
| CEP131 | Q9UPN4 | SMIM14 | Q96QK8 | WLS | Q5T9L3 |
| EPN1 | Q9Y6I3 | MAGED1 | Q9Y5V3 | MIB1 | Q86YT6 |
| STAM2 | O75886 | SCAMP1 | O15126 | SLC29A1 | Q99808 |
| ATP1A1 | P05023 | CALM3 | P0DP25 |  |  |
| ITGB1 | P05556 | HSPA8 | P11142 |  |  |

**Table S2B.**
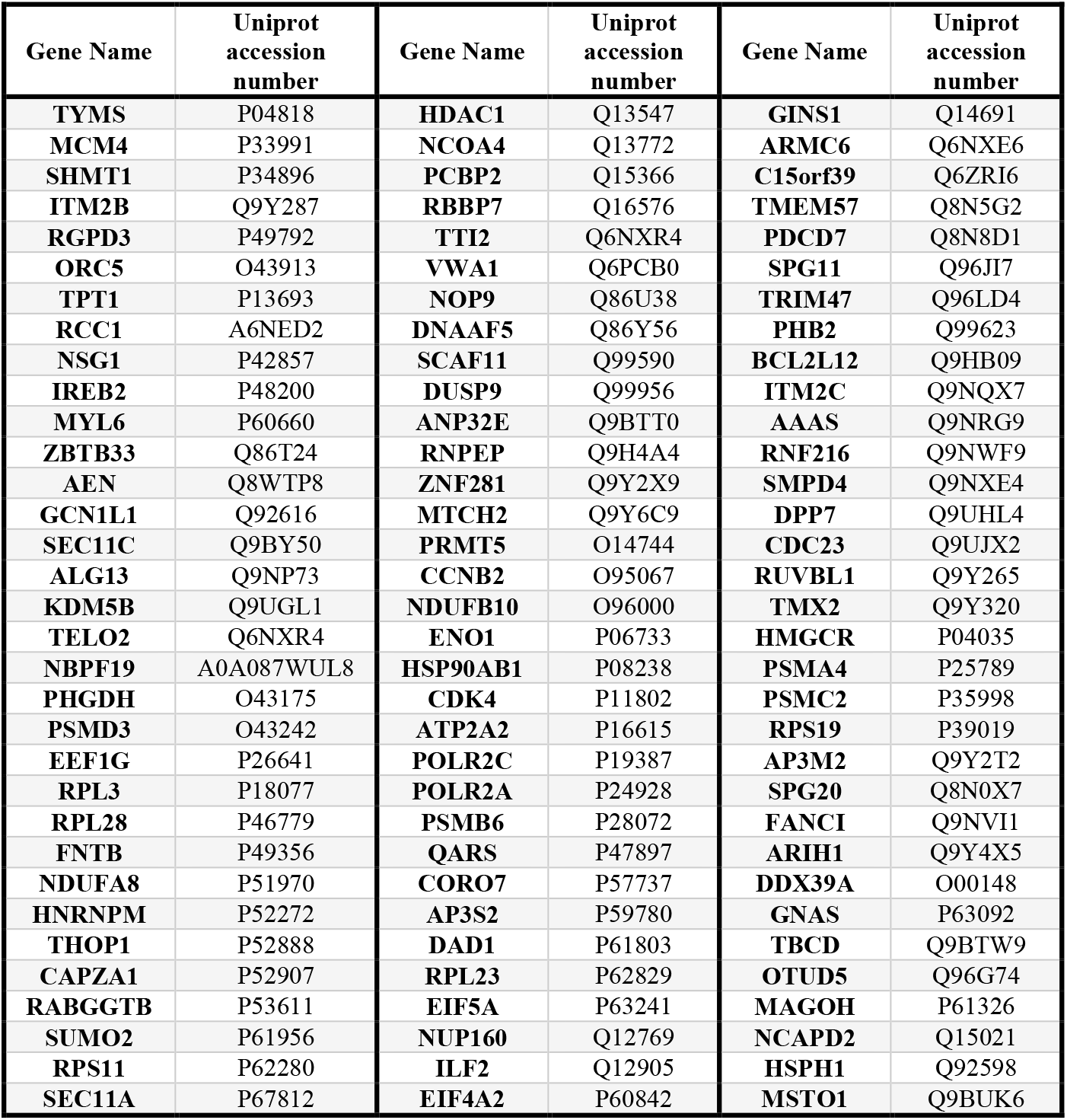
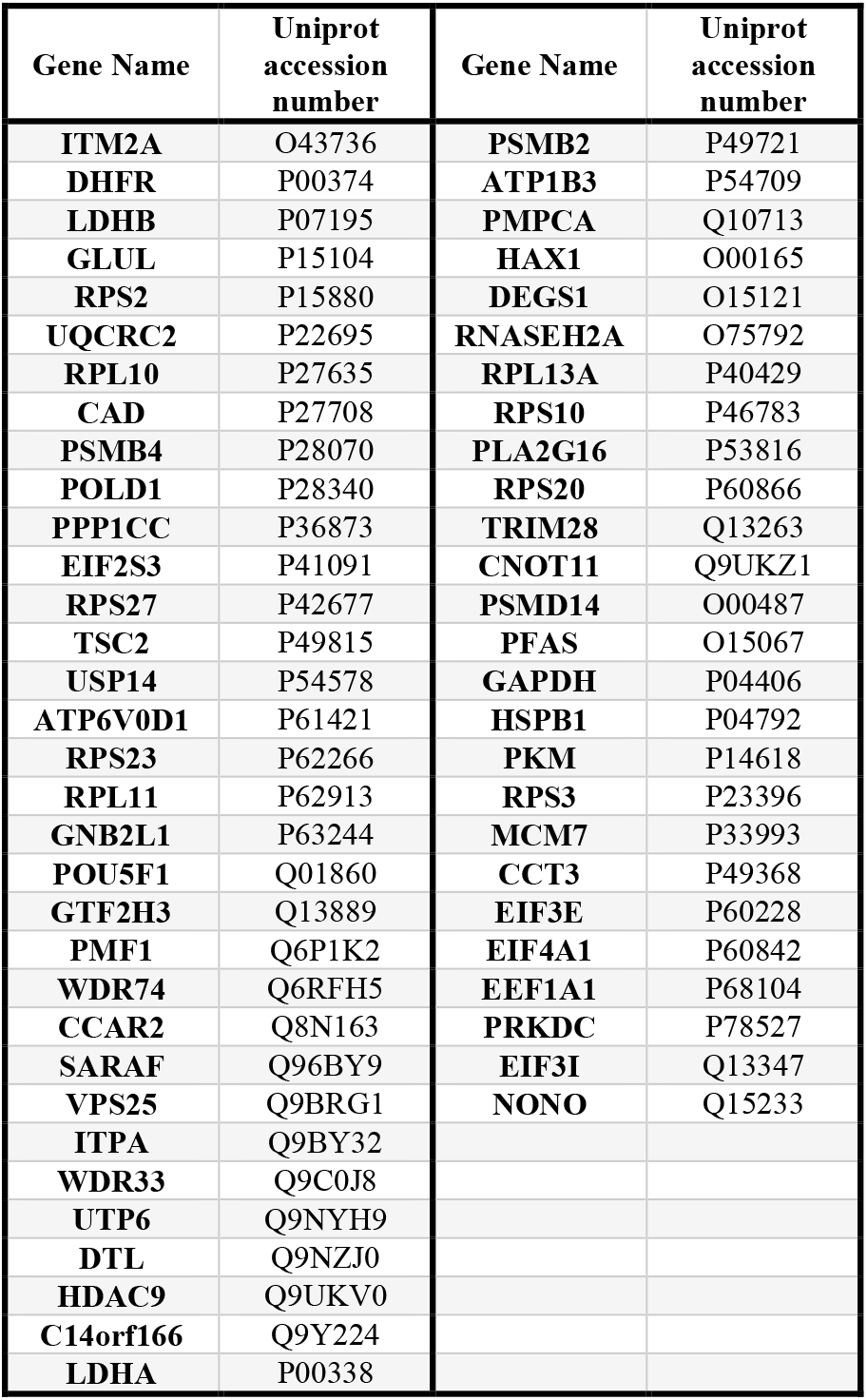
The table shows the differentially ubiquitinated proteins obtained in at least two out of four experiments, with log_2_ fold change ΔCSNAP / WT) below 0.67 in SILAC proteomic analysis

**Table S3.** Known CRL substrates identified in SILAC-based ubiquitination analysis of WT and ΔCSNAP cells. Based on references (1)(Emanuele et al., 2011), (2)^21^, (3)^22,^ and (4)^23^.

| Differentially ubiquitinated CRL substrates in WT and ΔCSNAP cells | | References | GO Annotation / Pathway<br>(Biological process / Molecular Function*) | | CRL substrate |
| --- | --- | --- | --- | --- | --- |
| **ΔCSNAP / WT down-regulated** | TYMS | 1 | DNA biosynthetic process | GO:0071897 |  |
|  | MCM4 | 1 | DNA replication | GO:0006260 |  |
|  | IREB2 | 1 | Metabolic process | GO:0008152 |  |
|  | KDM5B | 1 | Histone H3-K4 demethylation | GO:0034720 |  |
|  | PHGDH | 1 | Electron transport chain | GO:0022900 |  |
|  | PSMD3 | 1 | Ubiquitin-dependent protein catabolic process | GO:0006511 |  |
|  | RPL3 | 1,2 | Translation | GO:0006412 | CRL1 |
|  | HNRNPM | 1 | mRNA splicing, via spliceosome | GO:0000398 |  |
|  | ENO1 | 1 | Negative regulation of cell growth | GO:0030308 |  |
|  | POLR2C | 1 | Dual incision in TC-NER | R-HSA-6782135 |  |
|  | POLR2A | 1 | Dual incision in TC-NER | R-HSA-6782135 |  |
|  | RNF216 | 1 | Apoptotic process | GO:0006915 |  |
|  | PSMA4 | 1 | Proteasome | hsa03050+5685 |  |
|  | PSMC2 | 1 | Proteasome | hsa03050+5701 |  |
|  | ARIH1 | 1 | Protein polyubiquitination | GO:0000209 |  |
|  | OTUD5 | 1 | Protein deubiquitination | GO:0016579 |  |
|  | LDHB | 1 | Oxidation-reduction process | GO:0055114 |  |
|  | GLUL | 1 | Cell proliferation | GO:0008283 |  |
|  | UQCRC2 | 1 | Aerobic respiration | GO:0009060 |  |
|  | PPP1CC | 1 | Cell division | GO:0051301 |  |
|  | DTL | 1,2,4 | Cellular response to DNA damage stimulus | GO:0006974 | CRL1, CRL2/CRL4 |
|  | PSMB2 | 1 | Proteasome | hsa03050+5690 | CRL1 |
|  | TRIM28 | 1 | DNA repair | GO:0006281 |  |
|  | PFAS | 1 | Metabolic pathways | hsa01100+5198 |  |
|  | GAPDH | 1 | Metabolic pathways | hsa01100+2597 |  |
|  | RPS3 | 1 | Apoptotic process | GO:0006915 |  |
|  | MCM7 | 1,4 | Cell proliferation | GO:0008283 | CRL4,CRL2 |
|  | PRKDC | 1 | Cell proliferation | GO:0008283 |  |
|  | EIF3I | 1 | Translational initiation | GO:0006413 |  |
|  | NONO | 1,2 | DNA repair | GO:0006281 | CRL1 |
|  | CDC23 | 1 | Cell division | GO:0051301 | CRL3 |
|  | TPT1 | 2 | Cell differentiation | GO:0030154 | CRL1 |
|  | EEF1G | 2 | Eukaryotic Translation Elongation | R-HSA-156842 | CRL1 |
|  | CDK4 | 2 | Cell division | GO:0051301 | CRL1 |
|  | QARS | 2 | Metabolic pathways | hsa01100+5859 | CRL1 |

| Differentially ubiquitinated CRL substrates in WT and ΔCSNAP cells | | Reference<br>s | GO Annotation / Pathway<br>(Biological process / Molecular Function*) | | CRL<br>substrate |
| --- | --- | --- | --- | --- | --- |
| ΔCSNAP / WT<br>down-regulated | ITM2C | 4 | Positive regulation of extrinsic apoptotic signaling pathway | GO:2001238 | CRL2 |
|  | SMPD4 | 4 | Metabolic pathways | hsa01100+55627 | CRL2 |
|  | GNB2L1 | 3,4 | Apoptotic process<br>Cell cycle | GO:0006915<br>GO:0007049 | CRL1,<br>CRL2/CRL4 |
|  | HSP90AB1 | 1 | Negative regulation of proteasomal ubiquitin-dependent protein catabolic process | GO:0032435 |  |
|  | RPL13A | 1 | Translation | GO:0006412 |  |
|  | NCAPD2 | 4 | Cell division | GO:0051301 | CRL2 |
|  | AAAS | 4 | Nucleocytoplasmic transport | GO:0006913 | CRL4 |
|  | GNAS | 4 |  |  | CRL4 |
|  | DPP7 | 2 | Proteolysis | GO:0006508 | CRL1 |
|  | RPS2 | 2 | Translation | GO:0006412 | CRL1, CRL4 |
|  | HAX1 | 2 | Regulation of apoptotic process | GO:0042981 | CRL1 |
|  | RBBP7 | 4 | Cell proliferation | GO:0008283 | CRL2/CRL4 |
|  | CORO7 | 4 | Protein transport | GO:0015031 | CRL2/CRL4 |
|  | ARMC6 | 4 |  |  | CRL2 |
|  | PHB2 | 4 | Positive regulation of cell cycle G1/S phase transition | GO:1902808 | CRL2 |

\*GO Annotation Molecular Function

| Differentially ubiquitinated<br>CRL substrates in WT and<br>ΔCSNAP cells | | References | GO Annotation / Pathway<br>(Biological process) | | CRL substrate |
| --- | --- | --- | --- | --- | --- |
| **ΔCSNAP /**<br>**WT**<br>**up-**<br>**regulated** | PEX19 | 1 | Peroxisome | hsa04146+5824 |  |
|  | PPP2CA | 1 | Apoptotic process | GO:0006915 |  |
|  | EPN1 | 1 | Endocytosis | hsa04144+29924 |  |
|  | RPS3 | 1 | Apoptotic process<br>Cell division<br>DNA repair | GO:0006915<br>GO:0051301<br>GO:0006281 |  |
|  | HIF1A | 1,3 | Neddylation | R-HSA-8951664 | CRL1 |
|  | PI4K2A | 1 | Metabolic pathways | hsa01100+55361 |  |
|  | RAD23B | 1 | Nucleotide-excision<br>repair | GO:0006289 |  |
|  | HSPA8 | 1 | Regulation of cell<br>cycle | GO:0051726 |  |
|  | YBX1 | 1 | Positive regulation<br>of cell division | GO:0051781 |  |
|  | HIST1H1C | 1 |  |  |  |
|  | UBXN7 | 1 | Neddylation | R-HSA-8951664 |  |
|  | BCAP31 | 1 | Apoptotic cleavage<br>of cellular proteins | R-HSA-111465 |  |
|  | MIB1 | 1 | Protein<br>ubiquitination | GO:0016567 |  |
|  | RPN1 | 2 | Metabolic pathways | hsa01100+6184 | CRL1 |
|  | UBE2N | 4 | Positive regulation<br>of DNA repair | GO:0045739 | CRL2/CRL4 |
|  | MAGED1 | 3 | Regulation of<br>apoptotic process | GO:0042981 | CRL1 |
|  | RAD23A | 1 | Nucleotide-excision<br>repair | GO:0006289 |  |
|  | CALM3 | 2 |  |  | CRL1 |
|  | SLC6A8 | 4 | Transport | GO:0006810 | CRL4 |

**Table S4.**
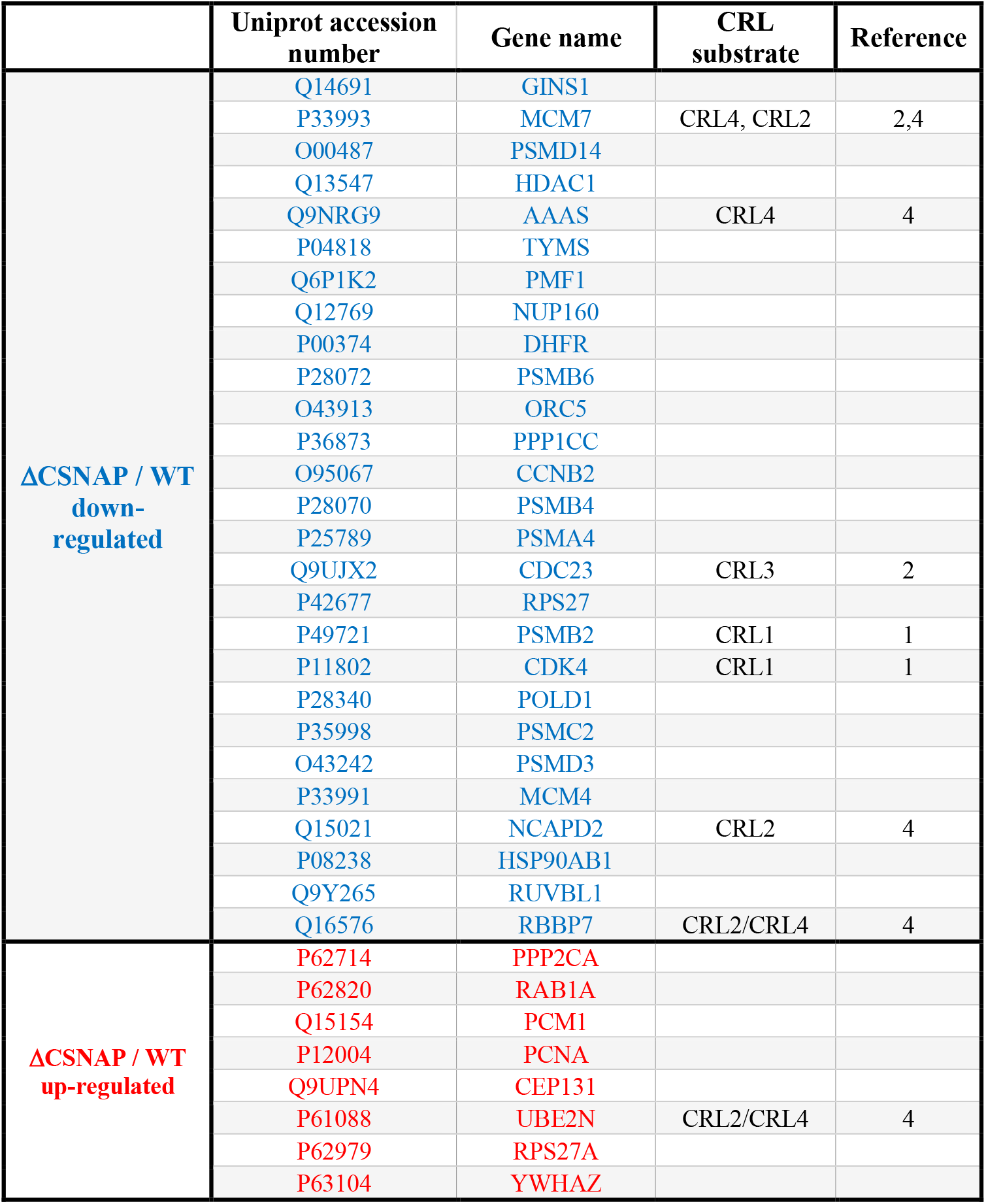
Differentially ubiquitinated proteins linked to cell cycle that were identified by SILAC-based analysis of WT and ΔCSNAP cells. Known CRL substrates are indicated based on the following references (1)(Emanuele et al., 2011), (2)^21^,, (3)^22^, and (4)^23^.

**Table S5.** Pathway analysis using Reactome 2016 of differentially ubiquitinylated proteins. Tables A and B display proteins exhibiting a ΔCSNAP/WT fold change bigger than 1.5 (A) or smaller then 0.66 (B).

The tables are provided as separate excel files.

**Table S6A.**
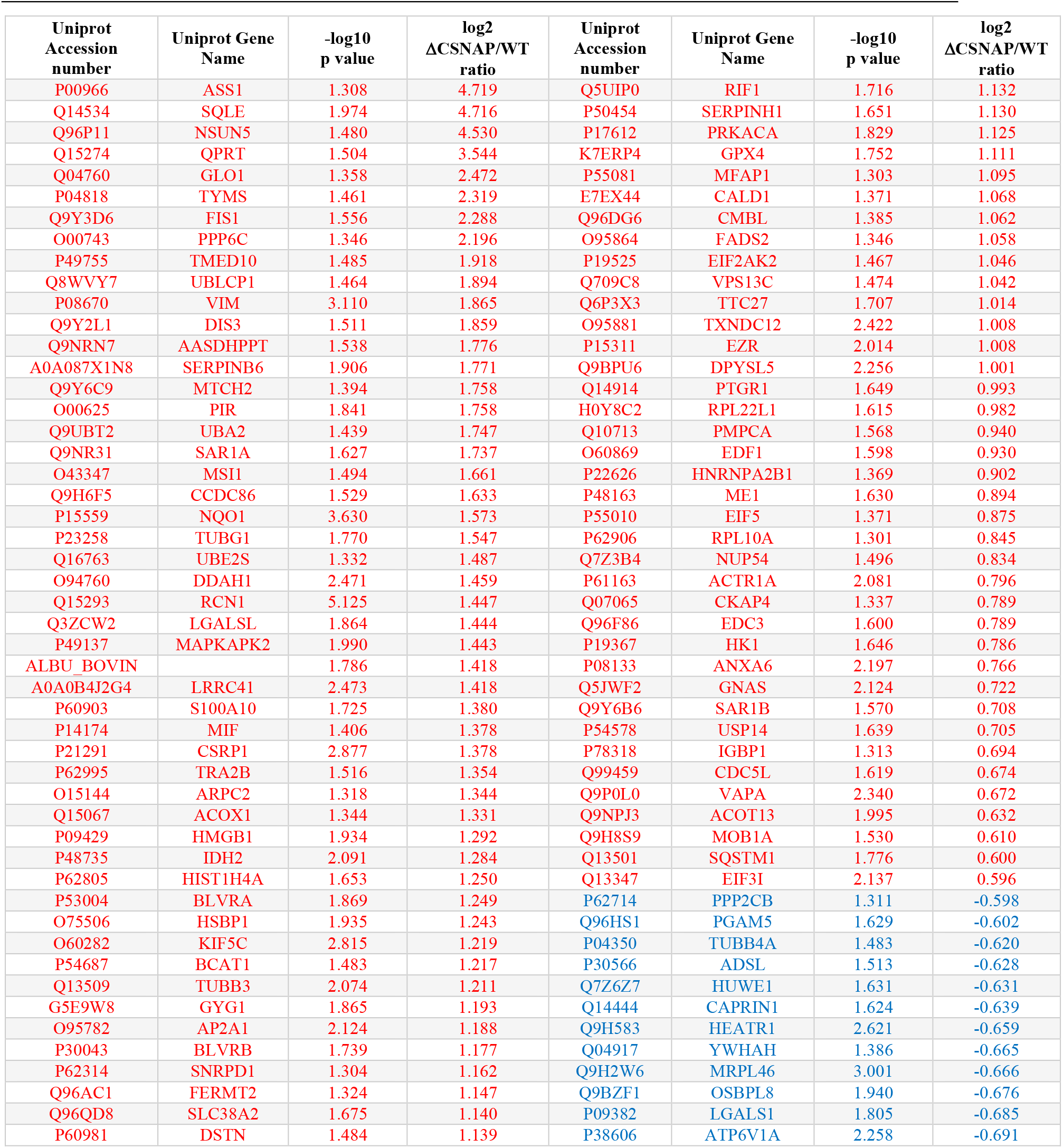
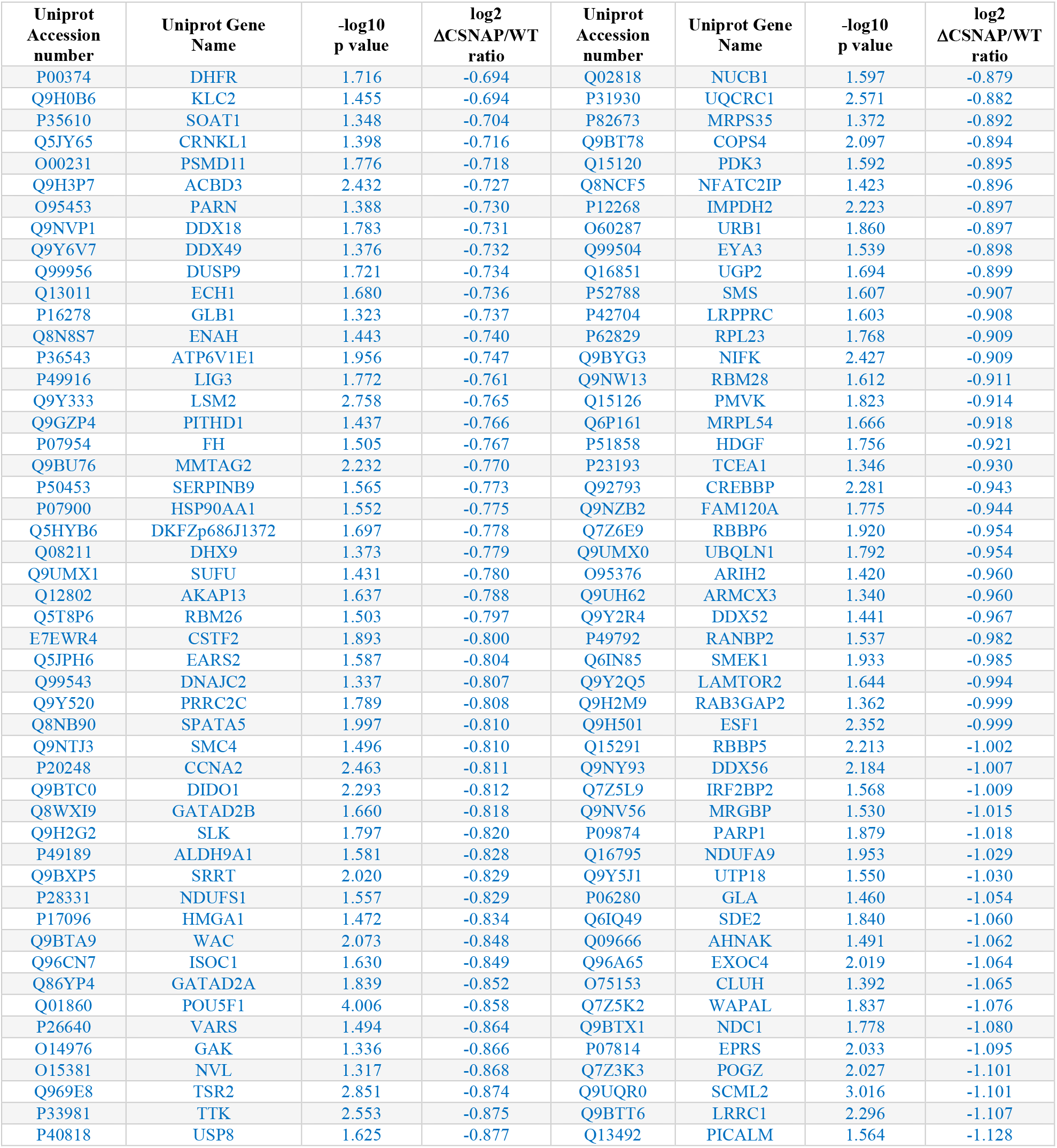
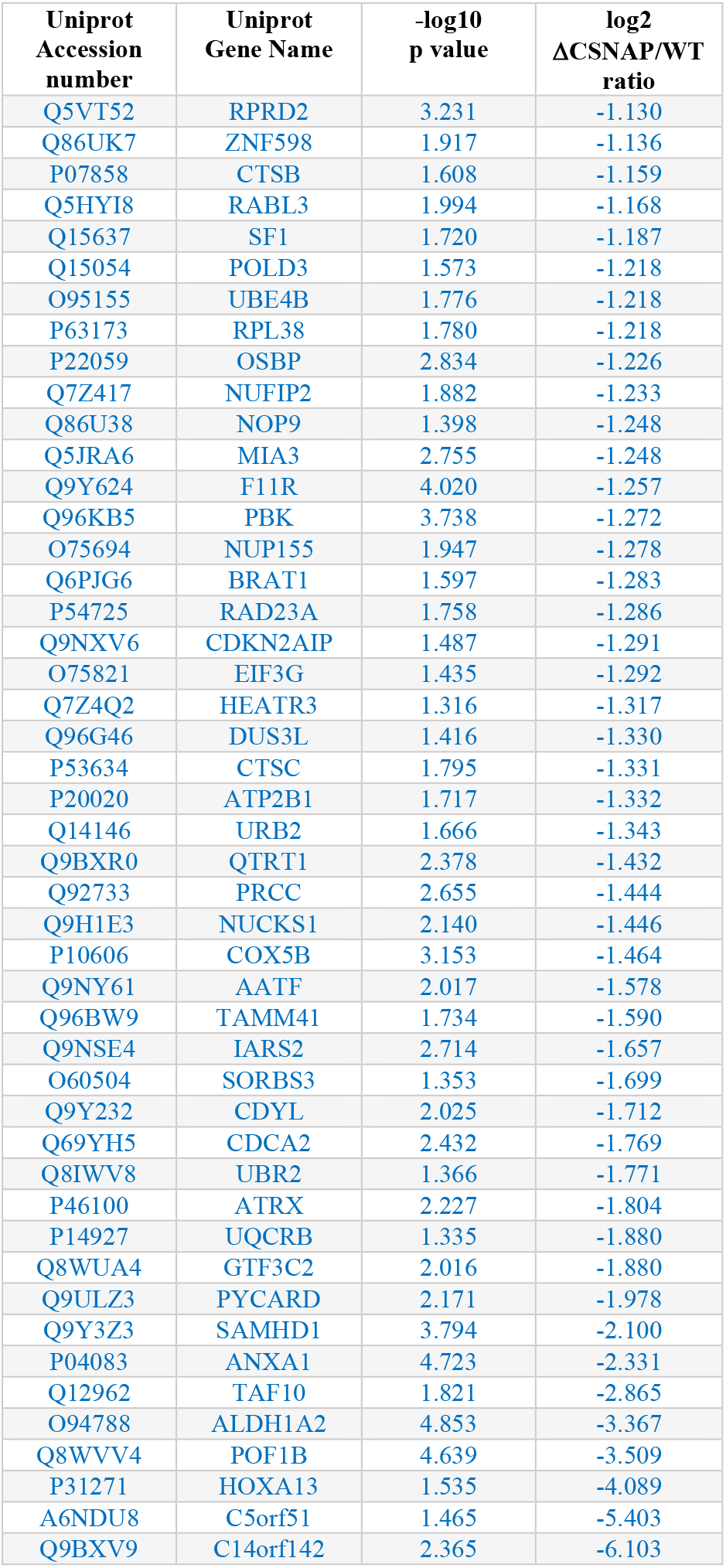
Proteins that were significantly differentially expressed between ΔCSNAP and WT cells, as demonstrated by total proteome label-free proteomic analysis. (-log_10_ p value>1.3; log_2_ fold changes x < −0.585 (blue) and x > 0.585 (red)).

**Table S6B.**
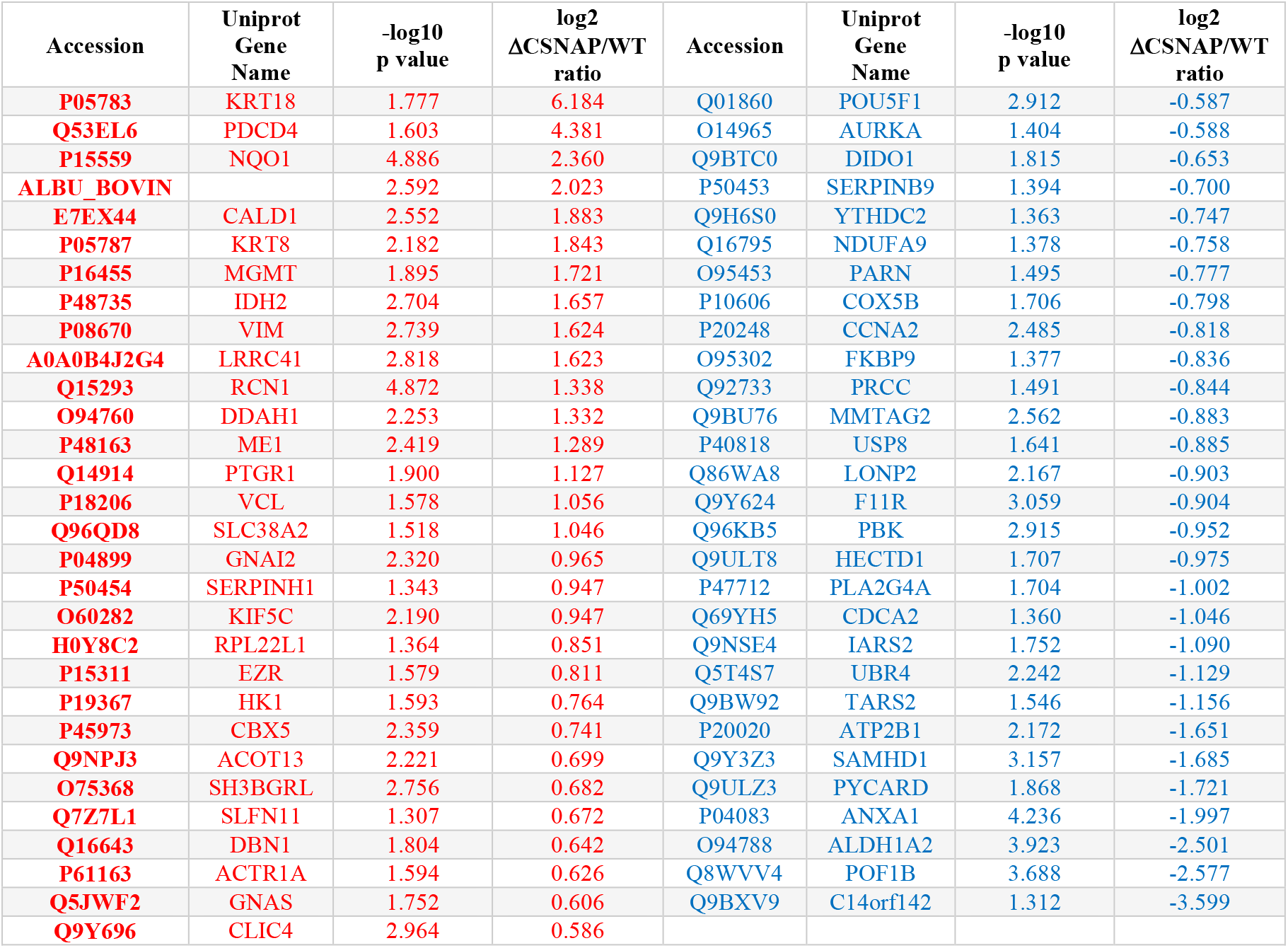
Proteins that were significantly differentially expressed between ΔCSNAP and WT cells following UV exposure, as demonstrated by total proteome label-free proteomic analysis. (-log_10_ p value>1.3; log_2_ fold changes x < −0.585 (blue) and x > 0.585 (red)).

**Table S6C.**
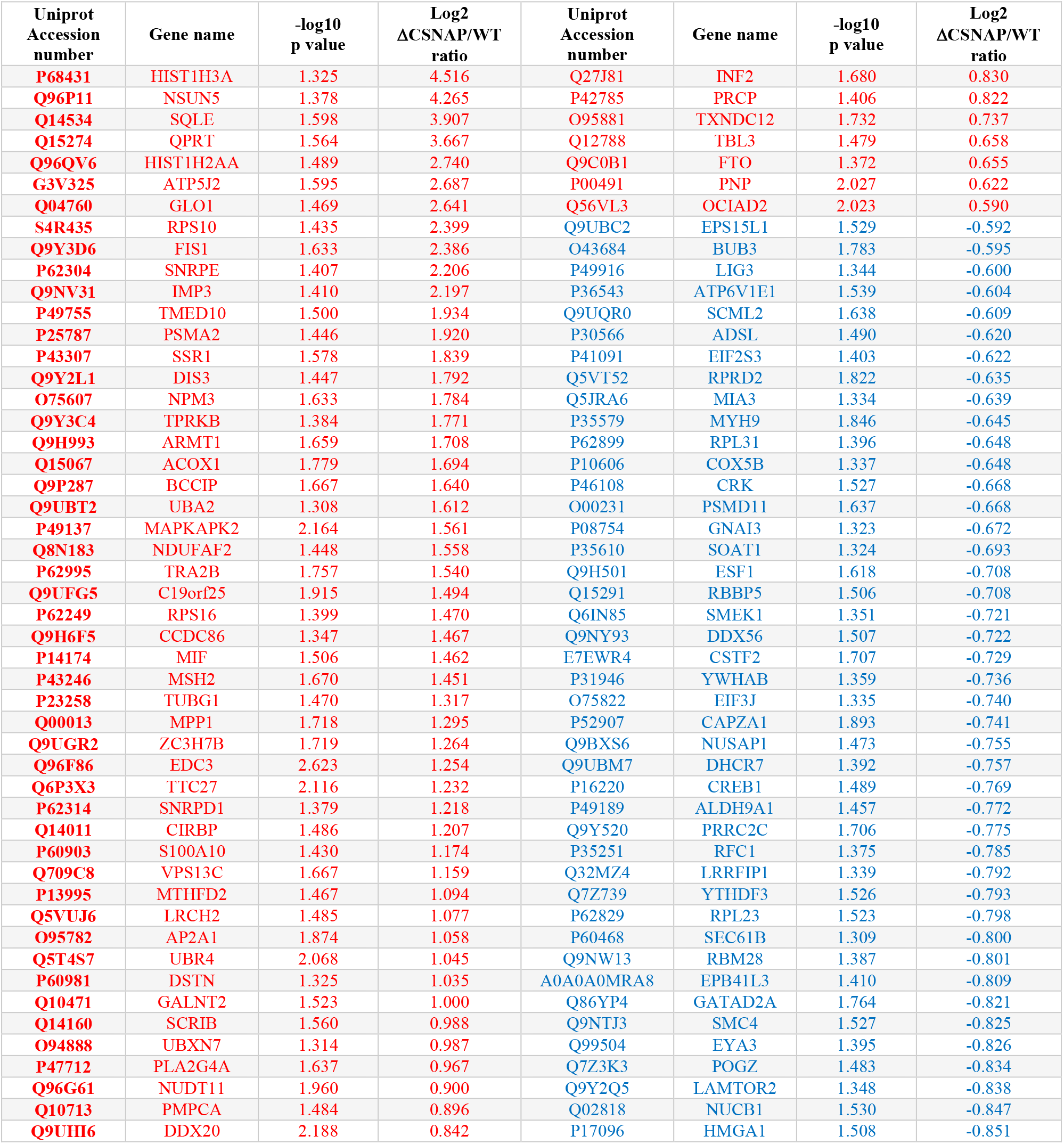
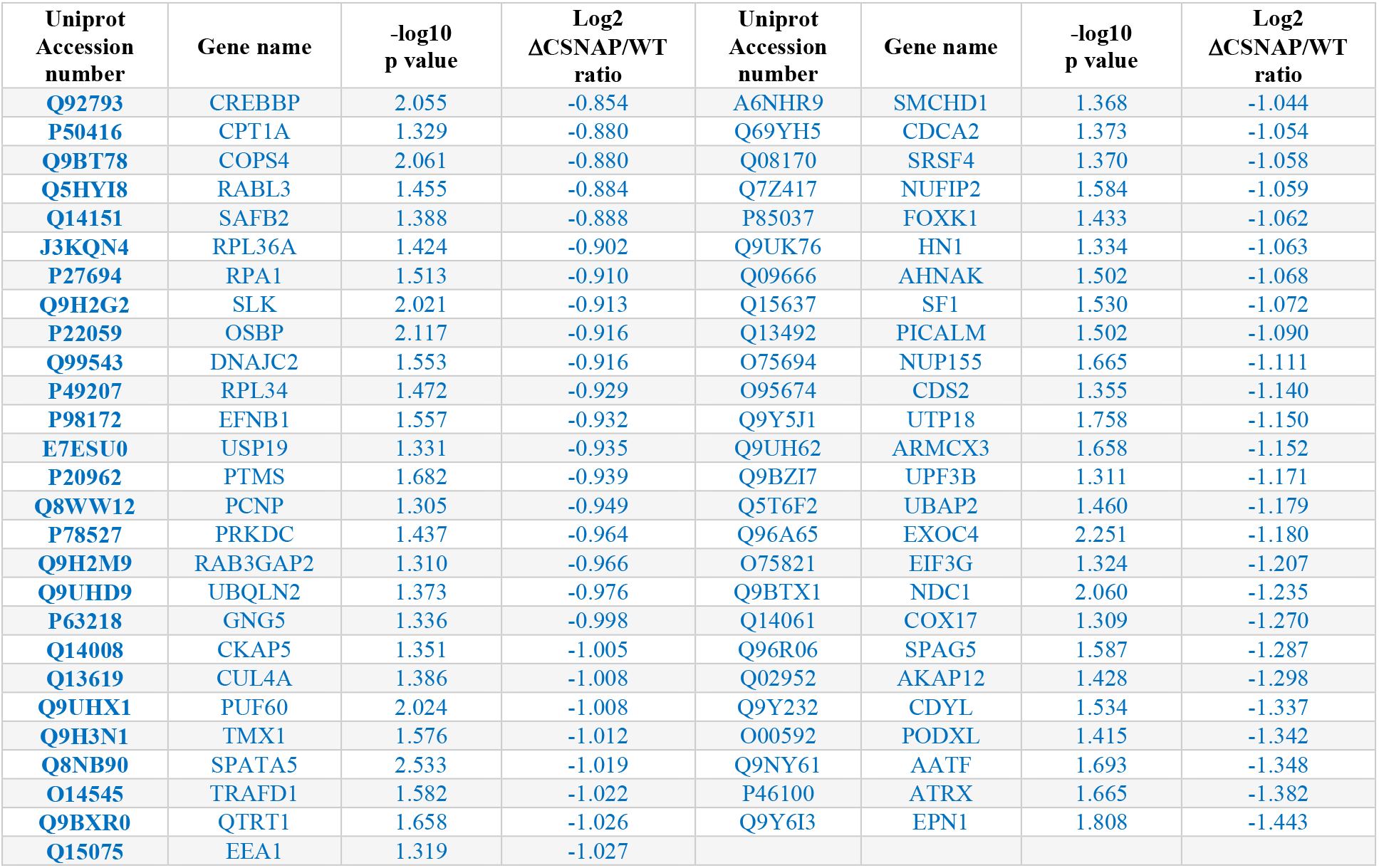
Proteins that were significantly differentially expressed between untreated and UV-exposed WT cells, as demonstrated by label-free proteomic analysis. (-log_10_ p value>1.3, log_2_ fold changes x < −0.585 (blue) and x > 0.585 (red)).

**Table S6D.** Proteins that were significantly differentially expressed between untreated and UV-exposed ΔCSNAP cells, as demonstrated by label-free proteomic analysis. (-log_10_ p value>1.3, log_2_ fold changes x < −0.58 (blue) and x > 0.58 (red)).

| Uniprot<br>Accession<br>number | Gene name | $-\log_{10}$<br>p value | $\log_2$<br>ΔCSNAP/WT<br>ratio |
| --- | --- | --- | --- |
| Q12962 | TAF10 | 1.344 | 2.203 |
| O94788 | ALDH1A2 | 1.493 | 0.933 |
| Q9BXR0 | QTRT1 | 1.336 | 0.853 |
| Q9UKJ3 | GPATCH8 | 1.384 | -1.005 |
| P20020 | ATP2B1 | 1.461 | -1.158 |

**Table S7.** Pathway enrichment analysis of whole cell label free proteomics data. The table is provided as a separate excel file.

**Table S8.**
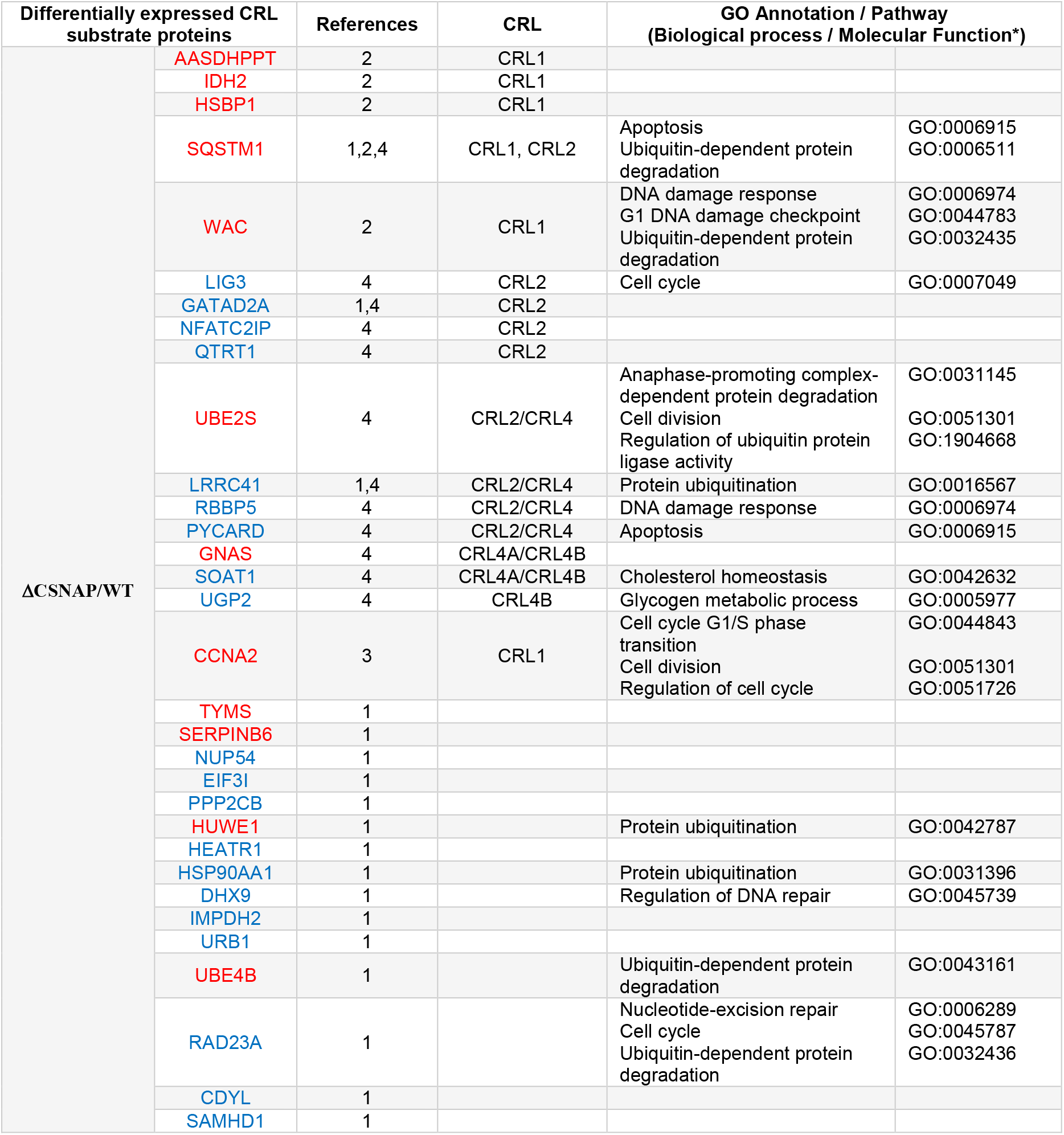
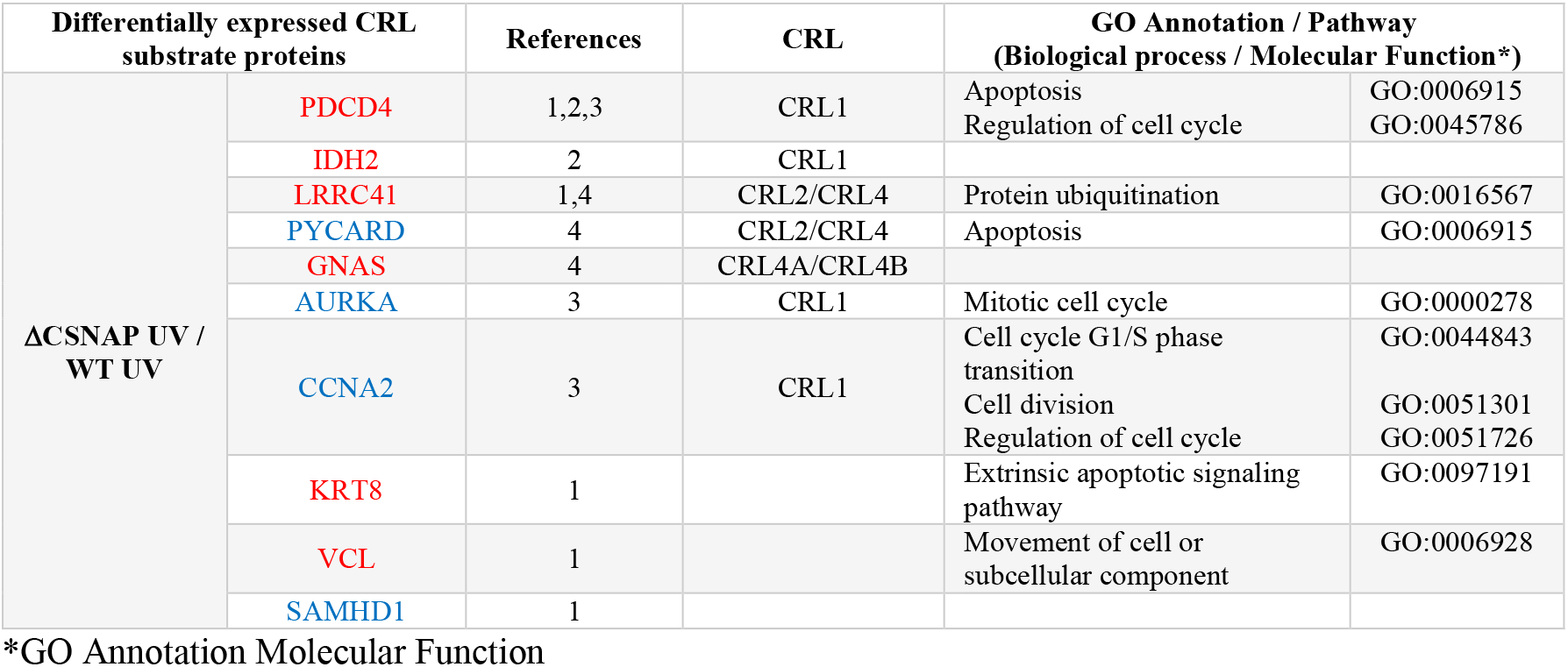
Known CRL substrates identified in total proteome analysis of WT and ΔCSNAP cells. Color coding represents up- (red) or downregulated (blue) proteins. Based on references (1)(Emanuele et al., 2011), (2)^21^, (3)^22^, and (4)^23^.

